# Differential modulation of Interferon and Cell Death Responses Define Human vs Avian Influenza A Virus Strain-Specific Virulence and guide Combination Therapy

**DOI:** 10.1101/2025.07.31.667854

**Authors:** Mansi Sharma, Abinaya Kaliappan, Dharmesh Bisoriya, Rajesh Thangavel Yadav, Rohan Narayan, Shachee Swaraj, Guojun Wang, Adolfo García-Sastre, Shashank Tripathi

## Abstract

Influenza A virus (IAV) poses a significant global health risk, with highly pathogenic strains like H5N1 (CFR ∼52%) causing severe disease compared to less lethal but more transmissible strains like H1N1 (CFR 0.01–0.03%). Although IAV primarily infects lung epithelial cells, causing cell death and tissue damage, the molecular basis of strain-specific pathogenesis remains poorly understood. Here we show that in cell culture, H5N1 induced more rapid and extensive cell death than H1N1. Since Interferon (IFN) signaling is key to innate immunity, we examined its role in virus-induced cell death using STAT1-knockout A549 cells and JAK/STAT pathway inhibitors like Baricitinib. Both approaches reduced cell death across various IAV strains, including H1N1, H5N1, H7N9, and H3N2. However, inhibition increased viral titers, raising concerns about its clinical use in isolation. To overcome this, we tested a combination of Oseltamivir (antiviral) and Baricitinib (anti-inflammatory). Post-infection treatment in a murine model reduced lung inflammation and improved survival. Given that both drugs are FDA-approved, this approach has strong translational potential for clinical IAV treatment.

## Introduction

Influenza A virus (IAV) is a respiratory virus recognized for inducing a range of symptoms from mild cold cough, fever, headache, etc., to severe illness like pneumonia and acute respiratory distress syndrome (ARDS) [1]. IAV infections constitute a significant global health concern, responsible for substantial morbidity and mortality. Seasonal IAV alone accounts for an estimated 290,000-65,000 respiratory deaths annually worldwide [2], with the potential for even graver consequences during pandemics. IAV is a member of the family *Orthomyxoviridae* and is an enveloped, negative-sense, single-stranded RNA virus having a segmented genome [3]. IAV infection is accompanied by lysis of the infected cell *in vitro* and *in vivo*. IAV primarily targets the airway epithelial cells, often causing cell death and tissue damage [4–7]. Several forms of programmed cell death (PCD) have been reported in the context of IAV infection; however, apoptosis is the most well-studied, followed by pyroptosis and necroptosis. Apoptosis is characterized by hallmarks such as DNA fragmentation, plasma membrane blebbing, exposure of phosphatidylserine (PS) on the cell surface, etc., yet the plasma membrane remains intact; thus, apoptosis is also considered as silent cell death or an anti-inflammatory process [8]. However, continued presence of uncleared apoptotic cells might result in rupture of the membrane and release of proinflammatory cytokines. On the other hand, in pyroptosis and necroptosis, the plasma membrane ruptures and cellular contents are released, making them proinflammatory cell death pathways [9]. While moderate levels of cell death and inflammation are beneficial in clearing out the virus, excessive amounts can trigger hypercytokinemia or cytokine storm [10, 11]. Highly pathogenic strains such as 1918 H1N1 and H5N1 IAVs are known to cause server disease primarily due to cytokine storm [12, 13]. Hence becomes imperative for us to study the relationship between IAV and cell death. Although IAV is known to influence cell death during its life cycle, the exact molecular mechanism remains elusive. Initially, the viral NS1 protein acts as an anti-apoptotic agent to prevent host cell death and to get time to complete viral replication. However, in later stages, cell death actually facilitates the spread of newly formed IAV particles to the neighbouring cells [14]. Viral protein PB1-F2 is known to be pro-apoptotic, and IAVs that contain this protein are often highly pathogenic [15]. Highly pathogenic avian IAV (HPAIV), including H5N1, H7N9, etc, cause severe symptoms in birds as well as in humans, in case of occasional spillover; however, effective human-to-human transmission has not been reported to date [16]. In contrast, H1N1 (2009 pandemic strain, H1N1pdm), highly transmissible but less pathogenic, exhibits milder pathology. Therefore, understanding the modulation of cell death by IAV on a strain-specific basis is essential. Previous studies have highlighted distinct pathogeneses among the various strains of IAV [17, 18]. With the increasing number of mammal-to-mammal transmission cases being reported for H5N1 [19, 20], the risk of spillover of an H5N1 IAV capable of efficient transmission within the human population persists, creating concerns of an imminent pandemic.

In this study, we have focused on the two strains that are known to be distinct in their pathophysiology and virulence: A/California/04/2009 (H1N1) – H1N1 and A/Viet Nam/1203/2004 (H5N1) – H5N1. Our cell death assays revealed higher levels of death by H5N1 as compared to H1N1 IAV *in vitro* conditions. However, interestingly, we observed a higher bystander cell death in H1N1 IAV-infected conditions, prompting us to look into the involvement of the interferon (IFN) signaling pathway. Interferons belong to the family of cytokines that are secreted in response to viral infections and act as the first line of defence in the form of the innate immune system [21]. IFN response begins with recognition of Pathogen Associated Molecular Patterns (PAMPs) by cellular Pattern Recognition Receptors (PRRs) like RIG-I, MDA5, TLRs, etc. The signal is then relayed to different transcription factors like IRF3, IRF7, NF-kB, etc, through specific kinases, leading to the production of different types of IFNs. IFNs are then detected by their receptors in an autocrine or paracrine manner. Downstream signaling gets activated via the Janus tyrosine kinase – Signal transducer and activator of transcription (JAK-STAT) pathway that eventually leads to the transcription of several interferon-stimulated genes (ISGs). These ISGs act as antivirals and key regulators of innate immunity and inflammation [22]. Therefore, as expected, cells lacking STAT1 showed a reduced cell death phenotype. A similar phenotype was observed when we used JAK inhibitors like Baricitinib and JAK Inhibitor I. Our findings indicate that Baricitinib alone leads to an increase in viral titers, making it unsuitable as a monotherapy for managing cytokine storm and lung inflammation in IAV-infected patients. To address this, we combined Baricitinib with Oseltamivir. According to previous studies, oseltamivir is most effective when administered within 48 ours of symptom onset [23, 24]. Its reduced efficacy at later stages is likely due to its limited ability to prevent lung damage and the development of ARDS [25]. Based on this, our study proposes that the combination of Baricitinib and Oseltamivir may offer a more effective strategy for treating severe IAV infections by minimizing the lung injury.

## Results

### H5N1 IAV displays accelerated replication kinetics and higher cell death response, compared to H1N1 IAV in human lung epithelial cells

Several studies have highlighted varying levels of pathology associated with H1N1 and H5N1 infections [17, 26, 27]. The viral strains used in this study are [A/California/04/2009 (H1N1) and A/Viet Nam/1203/2004 (H5N1)]. IAV infection typically results in compromising the alveolar barrier by disrupting the tight junctions of airway epithelial cells. [6, 28] leading to tissue damage. Therefore, we reasoned that the two viruses might exhibit distinct phenotypes when compared in the context of cell death. Firstly, we examined the kinetics of viral replication for both strains, in A549 lung adenocarcinoma cells, and found that the H5N1 virus replicated faster and to higher titers (Figure 1a) when compared to the H1N1 virus. However, at 72 HPI, the titers for both the viruses were almost similar, suggesting that H5N1 probably causes greater cell death, hence a sudden drop in viral titer. Viral-RNA (v-RNA) copy number was also quantified using qRT-PCR (Figure 1b). In line with the viral titers, cells infected with H5N1 consistently showed higher Matrix gene (M- gene copy numbers than those infected with H1N1 at all assessed time points, indicating a faster replication rate for H5N1. These findings suggest that H5N1 replicates more rapidly than H1N1. Consequently, we assessed the ability of both viruses to induce cell death in A549 cells microscopically. Our observations revealed that the H5N1 virus induced earlier and overall higher levels of cell death than the H1N1 virus (Figure 1c). Next, we compared the two viruses’ ability to cause cell death by checking the percent cell viability using the MTT assay (Figure 1d). At each time point, H1N1-infected cells were more viable compared to H5N1-infected infected. To support these findings, we used propidium iodide (PI) staining. PI is a fluorescent dye used to stain nucleic acids primarily in dead cells. Figure 1e shows that the H5N1 virus induces more cell death than the H1N1 virus. To assess the status of infected versus uninfected cells, we used GFP reporter viruses where the viral NS1 protein was fused with GFP protein [29]. GFP expression in cells indicates successful infection and GFP intensity signifies the replication. To ensure a clear observation, we infected the cells at a lower MOI (0.1), allowing sufficient presence of both dead and live cells. It was observed microscopically that the H5N1 GFP virus could significantly infect a greater number of cells and cause greater cell death when stained with PI compared to the H1N1 GFP virus (Figure 2a, b). Microscopic image analysis was conducted to quantify GFP-positive (infected) cells (Figure 2c), as well as cells that were double-positive for GFP and PI (infected and dead); and GFP-negative, PI-positive (bystander dead cells). These populations together accounted for the total PI-positive (dead) cells (Figure 2d). The analysis revealed that total cell death was significantly higher in cells infected with H5N1. Interestingly, however, a higher level of bystander (uninfected) cell death was observed in H1N1-infected samples compared to those infected with H5N1. These observations were corroborated by flow cytometry, which used dot plots to identify four cell populations: Q1 – GFP– PI+ (bystander dead cells); Q2 – GFP+ PI+ (infected and dead cells); Q3 – GFP+ PI– (infected and viable cells); Q4 – GFP– PI– (uninfected and viable cells) (Figure 2e). Over a time course of 12 to 72 hours post-infection (HPI), H5N1-infected cells consistently showed a higher proportion of GFP-positive cells (Figure 2f), increased overall cell death (Figure 2g), and reduced bystander cell death compared to H1N1-infected cells (Figure 2g). In summary, these findings suggest that H5N1 influenza A virus (IAV) is more virulent *in vitro*, leading to greater overall cell death, while H1N1 infection results in relatively higher bystander (uninfected) cell death.

**Figure 1:**
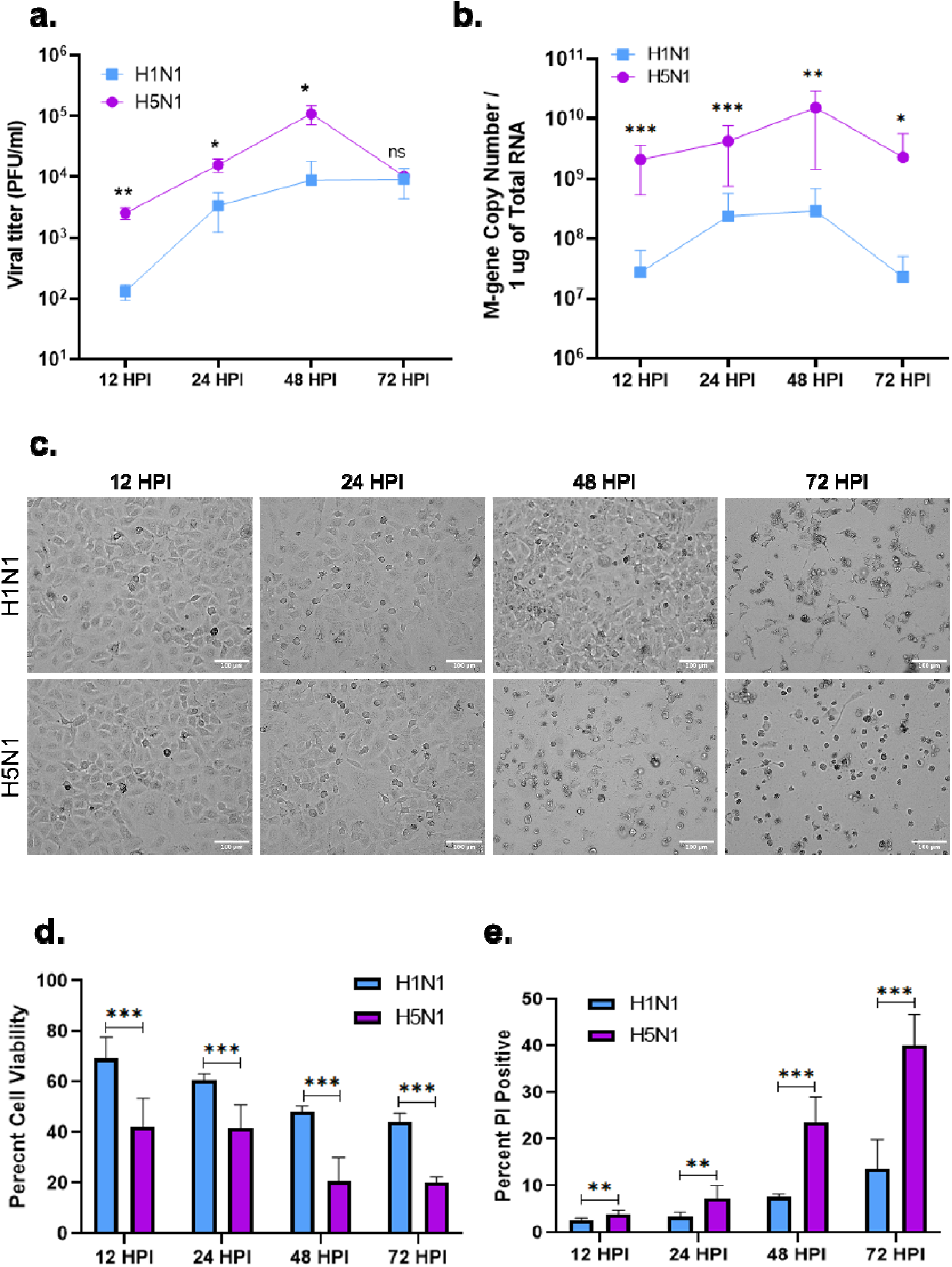
H5N1 Influenza A virus (IAV) replicates faster than H1N1 IAV and causes more cell death in human lung epithelial cells. **(a)** A549 cells were infected with either H1N1 or H5N1 virus at an MOI of 0.1 and viral titer was determined by plaque assay (**b**) and M-gene copy number was determined via qRT-PCR **(c)** A549 cells were infected with either H1N1 or H5N1 virus at 1 MOI and brightfield images were captured at 12, 24, 48 and 72 hours post infection (HPI) **(d)** A549 cells were infected with either H1N1 or H5N1 at 1 MOI and cell viability was determined using **(d)** MTT assay and **(e)** PI staining via flow cytometry. Data are presented as mean ± standard deviation (SD) from triplicate samples of a single experiment and are representative of results from three independent experiments. ***, p < 0.001, **, p < 0.01, and *, p < 0.05, by two-way ANOVA mixed effects model with Geisser-Greenhouse correction and Sidak’s multiple comparison test (a and b) and Multiple t-test (d and e).

**Figure 2:**
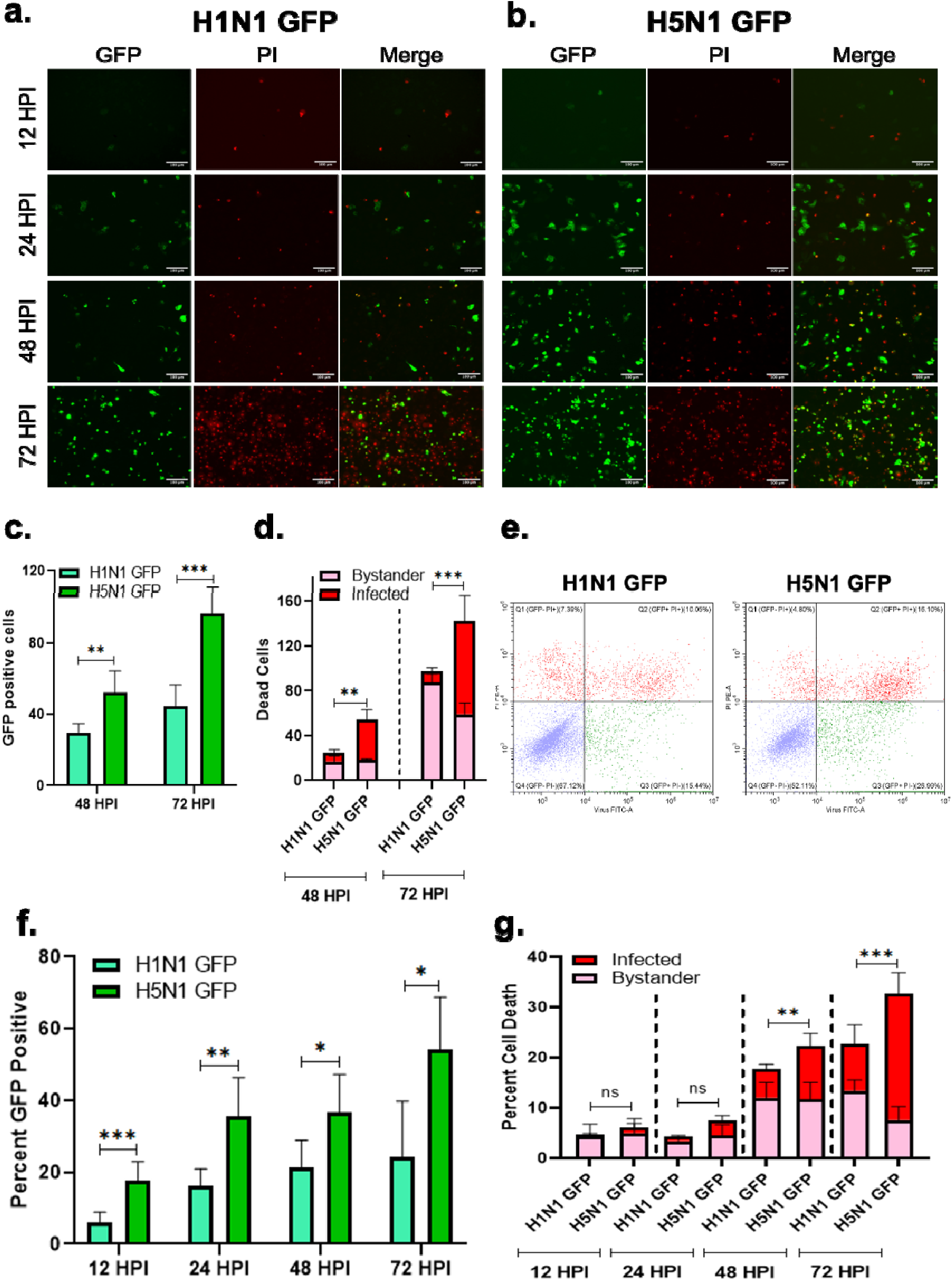
GFP-Reporter H5N1 and H1N1 IAVs highlight differences in infected vs bystander cell death induction. **(a)** A549 cells were infected with either H1N1 or **(b)** H5N1 GFP reporter virus at an MOI of 0.1 followed by staining of cells with PI and images were captured at 12, 24, 48 and 72 hours post infection (HPI). The images were analyzed to quantify: **(c)** GFP-positive (infected) cells; **(d)** cells that were both GFP+PI+ (infected and dead), as well as GFP–PI+ (bystander dead cells), which together represent the total number of dead (PI+) cells. Total GFP+, PI+, and GFP+PI+ cells were counted from six different fields and plotted as cell numbers. **(e)** Flow cytometry-based Dot plots showing the quadrants: Q1 – GFP- PI+ (Bystander dead cells); Q2 – GFP+ PI+ (Infected and dead cells); Q3 – GFP+ PI- (Infected and not dead cells); Q4 – GFP- PI- (Uninfected and not dead cells). **(f)** A549 cells were infected with either H1N1 or H5N1 GFP reporter virus at 0.1 MOI and percent GFP positive cells, **(g)** cells that were both GFP+PI+ (Q2 infected and dead), as well as GFP–PI+ (Q1 bystander dead cells), which together represent the total number of dead (PI+) cells were quantified using flow cytometry. Data are presented as mean ± standard deviation (SD) from triplicate samples of a single experiment and are representative of results from three independent experiments. ***, p < 0.001, **, p < 0.01, and *, p < 0.05, by two-way ANOVA mixed effects model with Geisser-Greenhouse correction and Sidak’s multiple comparison test. The statistics reflect Total cell death.

### H5N1 and H1N1 IAVs differentially modulate IFN response and cell death pathways in human lung epithelial cells

Given the observed differences in bystander cell death, we hypothesized that neighboring cells were being influenced by cytokines (particularly IFNs) from the infected cells. IFNs serve to alert the neighboring cells of impending viral infection [30] and are a key component of innate immunity that acts as an effective barrier for the virus to overcome to establish a productive infection. Given that IFN induction and signaling are crucial steps of our innate immune system to fight against viral infections, we sought to delve deeper into the differences in strain-specific cell death by examining the viruses’ ability to antagonize the interferon induction and signaling pathway. To this end, we looked into the activity of IFN-b and ISRE (IFN-stimulated response element) promoters during IAV infection. It was observed that H5N1 infection was unable to antagonize either of the two pathways (Figure 3a and b), suggesting that it may lack evolved mechanisms to counteract the human host’s immune response. Since cytokine release is among the strongest antiviral responses initiated by host cells, we examined the mRNA expression levels of various cytokines. We found that H5N1 infection led to a noticeable upregulation of many inflammatory cytokine genes (Figure 3c). Notably, the cells infected with H5N1 exhibited significantly higher IFN-β response compared to H1N1, consistent with previous reports [18, 31]. Other cytokines, including IFN- γ, λ, TNF-α, and proinflammatory cytokine IL-6 transcript levels were higher for cells infected with H5N1 compared to H1N1. The effects of IFN-λ are particularly prominent in epithelial cells, indicating its role in the specialized immune responses that safeguard epithelial surfaces, which are continuously exposed to respiratory viruses [32]. Therefore, there is a noticeably greater level of IFN-λ in cells infected with H5N1. Levels of antiviral genes such as IFIT2/ISG54 are also higher in the case of H5N1-infected cells. This could be explained by the overwhelming of the innate immune system by the rapid viral replication of the H5N1 virus compared to H1N1. Apoptosis, pyroptosis, and necroptosis are the three main programmed cell death pathways [33] involved during IAV infection. Hence, we examined the transcript levels of genes involved in these pathways during infection with either H1N1 or H5N1 IAVs (Figure 3d). Our results suggested that apoptosis-related genes (Caspase 8, Caspase 9, and BAX) were predominantly upregulated in H1N1-infected cells, whereas most of the pyroptosis (Caspase 1 and NLRP3) and necroptosis (MLKL) associated genes were more strongly upregulated in H5N1-infected cells. These findings suggest that H5N1 may induce more inflammatory cell death, potentially contributing to the more severe tissue damage observed. Whereas H1N1 IAV causes more apoptotic or silent cell death. To further validate our claims, we checked the expression levels of various cell death-related proteins (Figure 3e). Expression levels of cleaved caspase 3 and caspase 8 were higher in lysates from H1N1-infected cells, indicating a more apoptotic cell death (Figure 3f). However, the expression of cleaved Gasdermin-D (a downstream protein of the pyroptotic pathway) is higher for H5N1-infected cells, reconfirming the mode of cell death to be pyroptosis. Next, we utilized the publicly available FluOMICS database [34] to look closely into the differentially expressed genes (DEGs) following IAV infection. Transcriptomics data from Human tracheobronchial epithelial (HTBE) cells were analyzed for DEGs between cells infected with either H1N1 or H5N1 IAV. The gene cluster map showed distinct temporal gene expression patterns (Supplementary Figure 1a). Metascape analysis also pointed to differential expression of signaling pathways between the two viruses (Supplementary Figure 1b). Analysis revealed that pathways associated with antiviral defence, immune and interferon response were activated as early as 3 HPI for H5N1-infected cells, and the inflammatory response remained higher for H5N1 cells. This suggests that H5N1 infection elicits a stronger inflammatory response compared to H1N1 IAV. Within this dataset, inflammatory cytokines were specifically curated and compared between H1N1 and H5N1 (Supplementary Figure 1c). It was observed that most of the inflammatory cytokines were upregulated in the case of H5N1 compared to H1N1. Thus, we conclude that H5N1 and H1N1 IAVs differentially modulate IFN response and expression of cell death-related genes.

**Figure 3.**
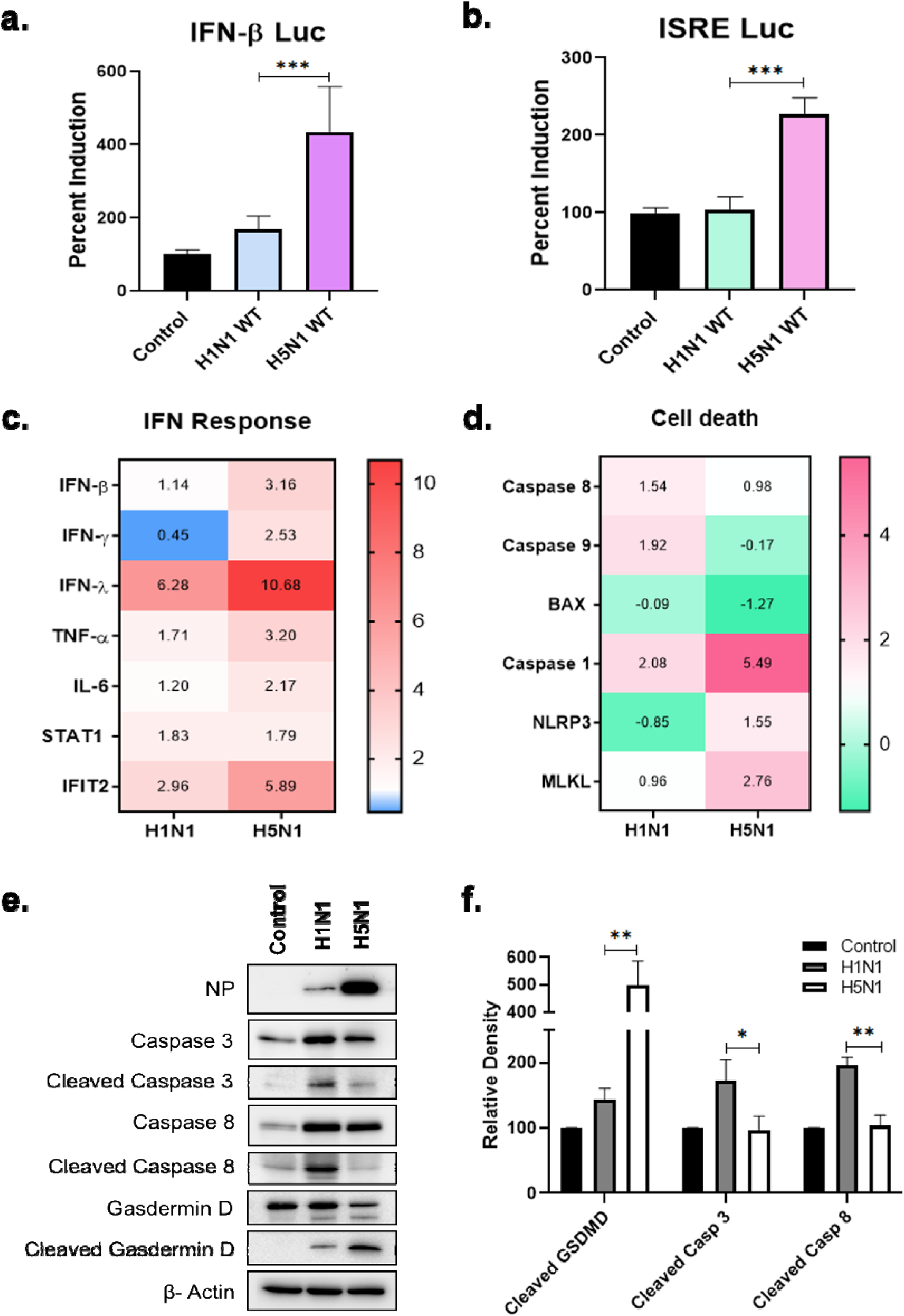
H5N1 and H1N1 IAVs differentially modulate IFN response and cell death pathways in human lung epithelial cells. **(a)** HEK293T cells were co-transfected with either IFN-β or **(b)** ISRE promoter-driven Firefly Luciferase reporter plasmid and Renilla Luciferase reporter plasmid. 24 hours post-transfection, cells were infected with either H1N1 or H5N1 IAV, for 16 hours, followed by assaying the cells for dual luciferase activity. **(c)** mRNA fold change (log_2_) levels of several IFN response genes and **(d)** host cell-death related genes were determined by qRT-PCR from total RNA isolated 48 HPI from A549 cells infected with either H1N1 or H5N1 WT IAVs. **(e)** Protein expression levels of viral NP, and several cell death markers including proactive and active forms of Caspase 3, 8 and Gasdermin D were observed via western blotting and **(f)** the band intensities were quantified. Data are presented as mean ± SD from triplicate samples of a single experiment and are representative of results from three independent experiments. ***, p < 0.001, **, p < 0.01, and *, p < 0.05, by Mann Whitney t-test (a and b) and Multiple t-test (f).

### STAT1-dependent IFN signalling promotes IAV-induced Cell death

The obvious involvement of IFN prompted us to explore the role of STAT1 signaling in the cell death phenotype seen during viral infection. We used STAT1 knockout (STAT1 KO) A549 cells, which were confirmed by western blotting (Supplementary Figure 2a). Expression of further downstream interferon-stimulated gene (ISGs) like ISG56/IFIT1 and ISG54/IFIT2 transcripts were also verified (Supplementary Figure 2b, c). These genes exhibited significant upregulation when the WT cells were treated with universal type-I IFN, whereas their gene expression was very low for STAT1 KO cells, indicating a defective IFN signaling pathway. Validated STAT1 KO and WT A549 were infected with either H1N1 or H5N1 GFP reporter virus, and 48 hours post-infection, cells were visualized via fluorescent microscopy (Figure 4a and b). Microscopic images were analyzed to quantify GFP-positive (infected) cells (Figure 4c), along with cells that were double-positive for GFP and PI (infected and dead), and GFP-negative, PI-positive (bystander dead cells). Together, these groups constituted the total PI-positive (dead) cell population (Figure 4d). A reduction in total and bystander cell death was observed for cells infected with either of the viruses. Due to the absence of a functional IFN signaling pathway, STAT1 KO infected cells showed increased GFP intensity, indicating enhanced viral replication, which was confirmed via flow cytometry as well (Figure 4c and e). As expected, reduced total cell death was evident in STAT1 KO cells for both viruses as observed through flow cytometry (Figure 4f), although to different extents (55.1% reduction in total cell death for cells infected with H1N1, while 42.9% for H5N1infected cells). Bystander cell death was also reduced in STAT1 KO cells for both viruses, however, with a greater extent for H1N1-infected cells. The greater reduction in cell death observed in H1N1-infected STAT1 KO cells may be attributed to the inherently lower IFN production in H1N1 infection, combined with the further impairment of the IFN signaling pathway in the absence of STAT1. Collectively, our findings suggest that STAT1- dependent IFN signaling contributes to a varied cell death response and potentially influences the pathology of the two viruses.

**Figure 4.**
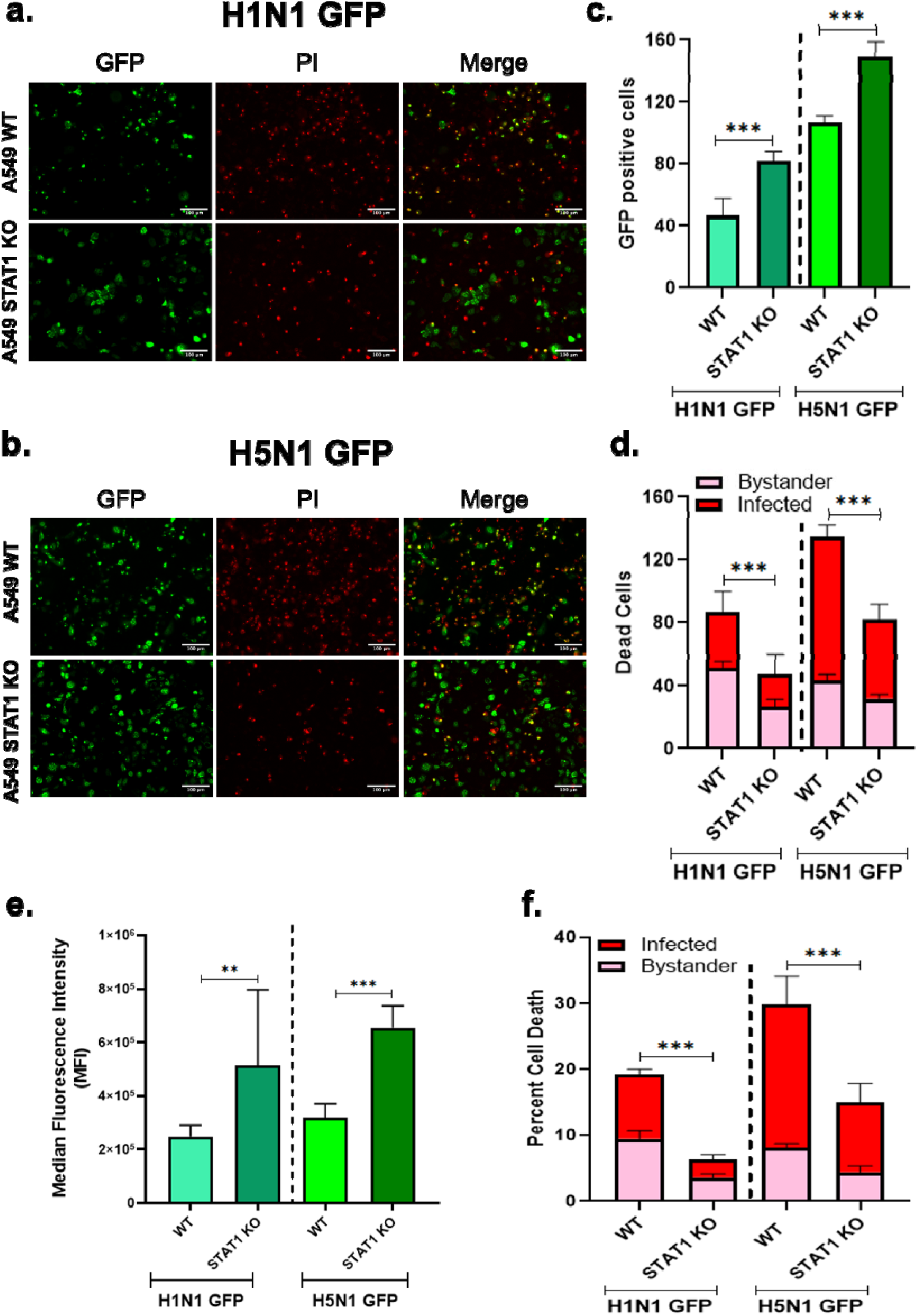
STAT1-dependent IFN signaling promotes IAV-induced cell death. **(a)** A549-WT and A549-STAT1 Knockout cells were infected with either H1N1 or **(b)** H5N1 GFP reporter virus at 0.1 MOI, fluorescent images were captured at 48 HPI and quantified as **(c)** GFP-positive (infected) cells; **(d)** cells that were both GFP+PI+ (infected and dead), as well as GFP–PI+ (bystander dead cells), which together represent the total number of dead (PI+) cells. Total GFP+, PI+, and GFP+PI+ cells were counted from eight different fields and plotted as cell numbers. **(e)** A549-WT and A549-STAT1 Knockout cells were infected with 0.1 MOI of either H1N1 or H5N1 GFP virus, Median Fluorescence Intensity MFI (viral replication), **(f)** cells that were both GFP+PI+ (Q2 infected and dead), as well as GFP–PI+ (Q1 bystander dead cells), which together represent the total number of dead (PI+) cells were quantified using flow cytometry. Data are presented as mean ± SD from triplicate samples of a single experiment and are representative of results from three independent experiments. ***, p < 0.001, **, p < 0.01, and *, p < 0.05, by Mann-Whitney t-test. The statistics reflect Total cell death.

### Chemical inhibition of STAT1-mediated IFN signaling ameliorates IAV-induced cell death

Upon sensing interferon (IFN) through its receptor, the associated kinase JAK becomes activated and phosphorylates STAT1. This initiates downstream signaling that culminates in the transcription of interferon-stimulated genes (ISGs). However, when JAK inhibitors are used, they block JAK activation, thereby preventing STAT1 phosphorylation and subsequently inhibiting the downstream IFN signaling cascade. Thus to strengthen our claims, we used two chemical inhibitors of IFN signaling, (i) Baricitinib which is an FDA approved drug for the treatment of rheumatoid arthritis, it is a selective and orally bioavailable JAK1 and JAK2 inhibitor and (ii) JAK Inhibitor I which is a potent, cell permeable ATP competitive inhibitor of JAK1, 2,3 and TYK2 [35, 36]. Baricitinib is an anti-inflammatory drug and recently repurposed for the treatment of SARS-CoV-2 in combination with remdesivir [37]. Cytotoxicity was checked for both the drugs, where CC_90_ was 4.3 and 4.8 μM for Baricitinib and JAK Inhibitor I, respectively (Supplementary Figure 3a). Further, both inhibitors were capable of blocking the phosphorylation of STAT1 (Supplementary Figure 3b), hence stalling the downstream STAT1 signaling. A549 WT cells were infected with either H1N1 or H5N1 WT or GFP reporter virus and 6 HPI cells were treated with 1 μM of either Baricitinib or JAK inhibitor I. 48 HPI, cells were stained with PI and visualized through microscopy and cell death was quantified using flow cytometry. Treatment with both inhibitors led to a reduction in virus-induced cell death during H1N1 and H5N1 WT infections, as observed through both microscopy and flow cytometry (Figure 5a and b). Similar effects were observed using the GFP reporter virus—treatment with JAK inhibitors resulted in enhanced viral replication, evidenced by increased GFP fluorescence intensity (Supplementary Figure 4a, b, and c) and elevated viral titers (Figure 5c). Cell death was assessed by quantifying GFP+PI+ (infected and dead) and GFP–PI+ (bystander dead) populations, which together represent the total PI-positive (dead) cells. Both total and bystander cell death were reduced in inhibitor-treated samples, with Baricitinib showing greater effectiveness (Supplementary Figure 4d and f). Consequently, subsequent investigations were carried out using Baricitinib. This observation was further validated by an increase in viral NP protein (Supplementary Figure 3b) in consistent with the increase in GFP levels in STAT1 KO cells and Baricitinib-treated cells. We also checked the transcript levels of various cytokines, including IFN-β, γ, and λ along with TNF-α, IL-6 and STAT1 and IFIT2 (ISG). In the presence of Baricitinib, a reduction in all cytokines was observed (Figure 5d). Furthermore, the transcript levels of cell death-related genes were also lower for the cells treated with Baricitinib (Figure 5e). Thus, we can conclude that chemical inhibition of IFN signaling reduces cell death in IAV-infected cells, probably via reduction in inflammatory cytokines. These results were further extended to different subtypes of IAV, including avian H7N9, H5N1, and human H3N2. There, too, we could observe a significant reduction in cell death in infected cells that were treated with Baricitinib (Figure 5f), while the increase in viral titer was determined by plaque assay (Figure 5g). This showed that the reduction in cell death due to inhibition of IFN-signaling is a universal phenomenon. Thus, we can conclude that inhibition of IFN-signaling reduces cell death; however, it increases the viral titers.

**Figure 5.**
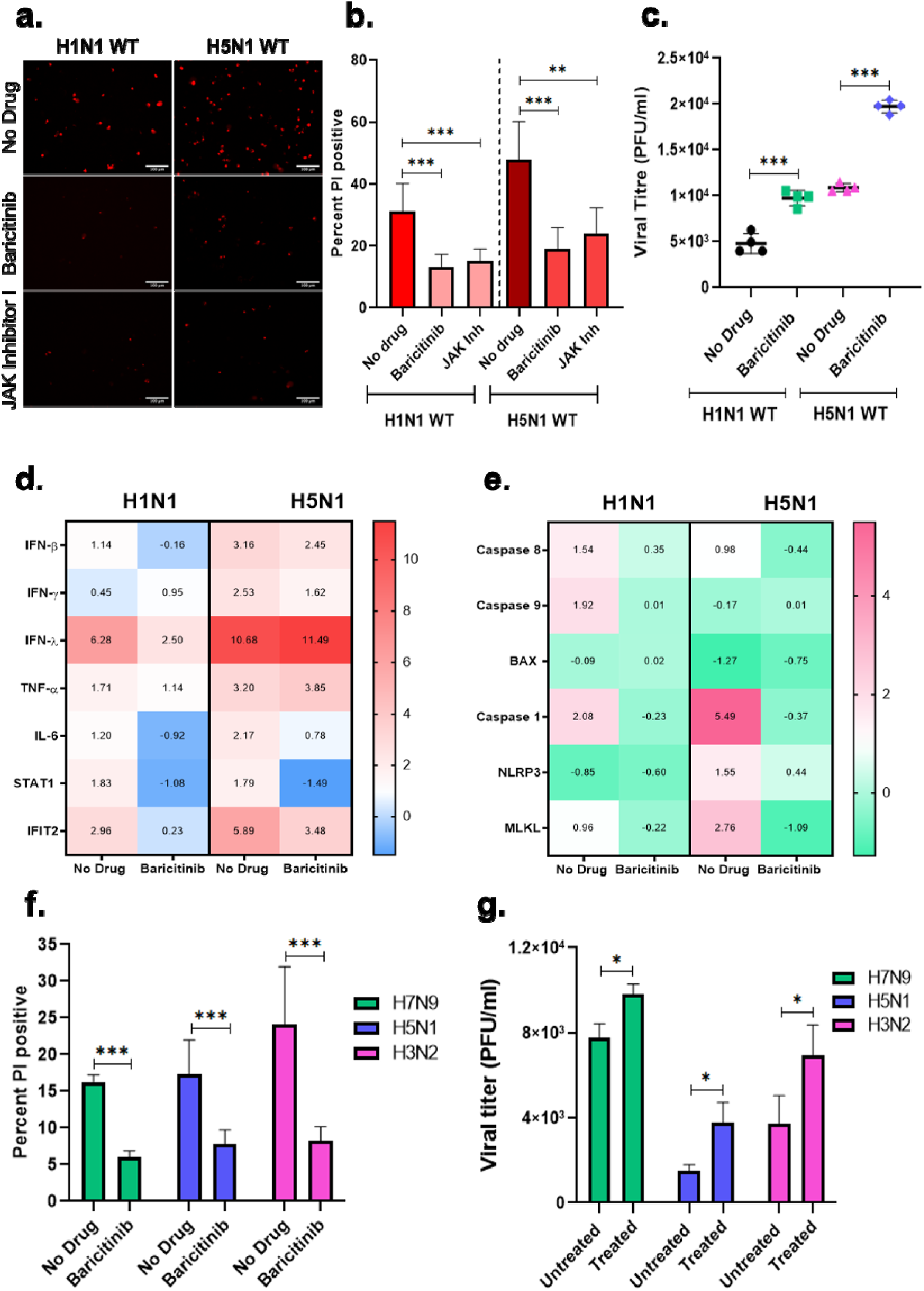
Chemical inhibition of STAT1 signaling ameliorates human respiratory epithelial cell death induced by a wide range of human and avian IAVs. **(a)** A549 cells were infected with either H1N1 or H5N1 WT virus at an MOI of 0.1. 6 HPI cells were treated with 1 µM either Baricitinib or JAK inhibitor I for 48 HPI, and subsequently total cell death was observed by **(a)** fluorescent microscopy and **(b)** quantified by flow cytometry while **(c)** viral titer of supernatants from untreated and Baricitinib treat cells was quantified by plaque assay. **(d)** mRNA fold change (log_2_) levels of several IFN response genes and **(e)** host cell-death related genes were determined by qRT-PCR from total RNA isolated 48 HPI from A549 cells infected with either H1N1 or H5N1 WT IAVs and treated with Baricitinib 6 HPI. **(f)** A549 cells were infected with avain H7N9, H5N1 or human H3N2 IAVs at an MOI of 0.1. 6 HPI cells were treated with 1 µM either Baricitinib for 48 HPI, and subsequently total cell death was quantified by flow cytometry and **(g)** the viral titer was estimated via plaque assay. Data are presented as mean ± SD from triplicate samples of a single experiment and are representative of results from three independent experiments. ***, p < 0.001, **, p < 0.01, and *, p < 0.05, by Brown-Forsythe and Welch ANOVA testes, Holm-Sidak’s multiple comparisons test (b) and Unpaired t-test with Welch’s correction (c, f and g).

### Combination of Baricitinib and Oseltamivir demonstrates superior anti-IAV antiviral efficacy, compared to standalone treatment in cell culture

Treatment of cells with Baricitinib alone led to increased viral titers, highlighting a potential risk if used in isolation in clinical settings. To address this, we explored a combination therapy involving Oseltamivir (a direct-acting antiviral) and Baricitinib (an anti-inflammatory drug). Oseltamivir, commonly known as Tamiflu, is a popular antiviral against the influenza virus [38]. It is a direct-acting antiviral and inhibits the activity of viral Neuraminidase (Na) protein, hence preventing the release of the virus from infected cells. As shown previously, inhibition of STAT1 signaling mitigated cell death and inflammation; however, it also led to an increase in viral infection. Therefore, we reasoned that a combination of the two drugs Baricitinib and Oseltamivir would be able to reduce the cell death as well as infection. Studies have shown that the combination of Baricitinib and Remdesivir is effective for hospitalized patients with SARS-CoV-2 infection [37]. To this end, we first assessed the cytotoxicity of Oseltamivir, determining the CC_90_ to be 8.7 μM (Supplementary Figure 5a). Based on different combinations tried (data not shown), using half the concentration of Baricitinib, which had been used earlier (i.e., 0.5 μM), and 1 μM of Oseltamivir, in combination, was used to check for the cytotoxicity of the combination. There was no significant difference observed in the cell viability between treatments with Baricitinib alone, Oseltamivir alone, or their combination (Supplementary Figure 5b). We next tested the combination in cells infected with either H1N1 or H5N1 WT IAVs. As expected, Baricitinib alone reduced cell death in the case of cells infected with either H1N1 or H5N1, while no significant difference was observed in cells treated with Oseltamivir alone. However, a significant reduction in cell death was observed when the combination was used (Figure 6a). Additionally, while Baricitinib treatment alone led to an increase in viral titers, Oseltamivir treatment resulted in a substantial reduction. Interestingly, the combination treatment significantly lowered viral titers (Figure 6b). Transcript levels of cytokines and cell death– related genes were assessed from infected cells following combination treatment, and it was observed that the majority exhibited reduced expression compared to untreated cells (Figure 6c and d). Further, combination treatment was also administered at a later time point (12 hours post-infection). Here, too, we observed a significant reduction in cell death (Figure 6e). This indicates that a delayed therapeutic intervention remains effective in reducing cell death and potentially mitigating infection-induced cell death. Thus, we can conclude that a combination of Baricitinib and Oseltamivir can act as an effective therapeutic against IAV infections.

**Figure 6:**
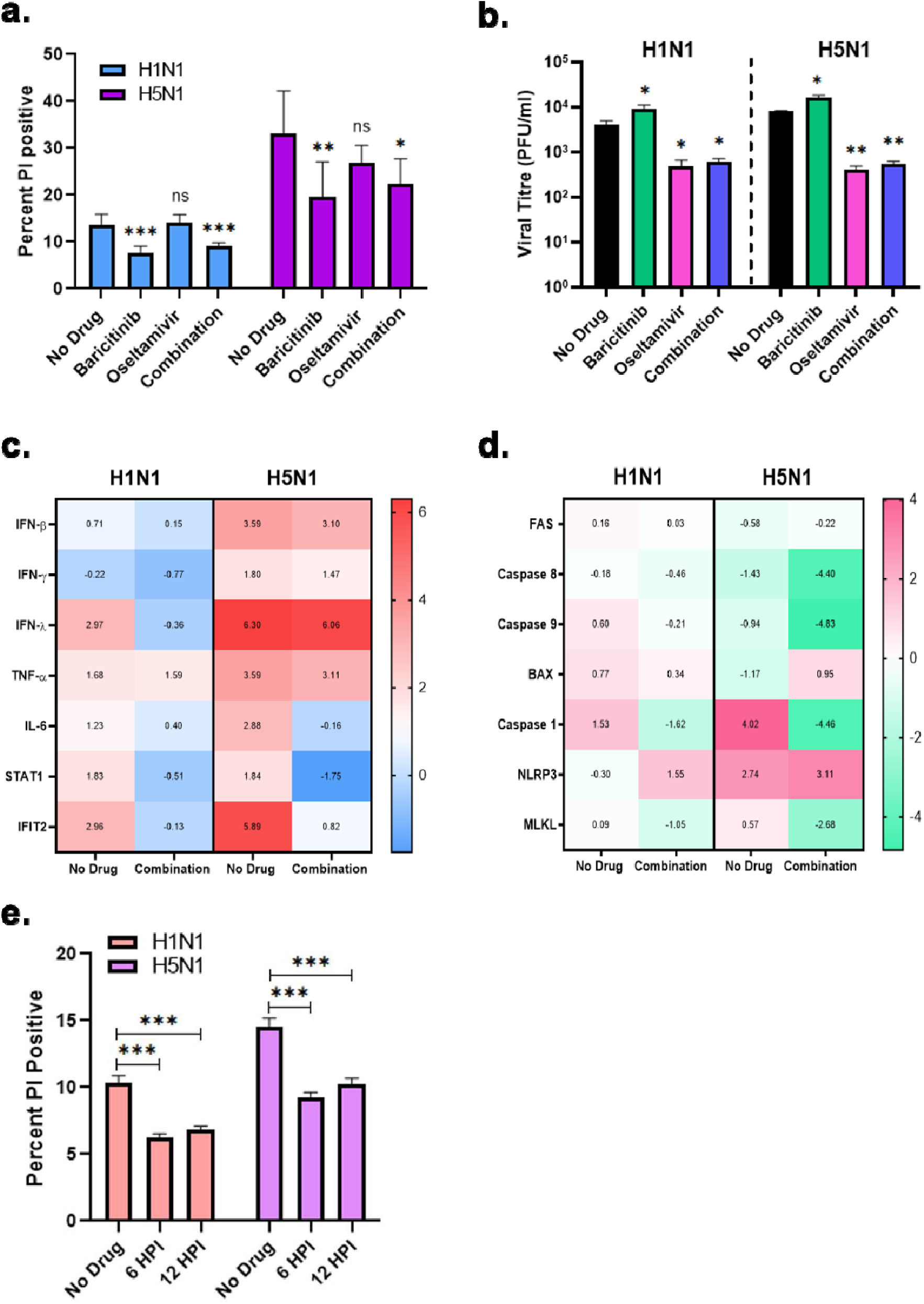
Combination of Baricitinib and Oseltamivir provides superior restriction of both IAV-induced cell death and viral replication. **(a)** A549 cells were infected with 0.1 MOI of either H1N1 or H5N1 WT virus followed by treatment with 1 µM of either Baricitinib, Oseltamivir alone or their combination (0.5µm of Baricitinib + 1µm of Oseltamivir) 6 HPI, subsequently 48 HPI total dead cells were quantified using flow cytometry and **(b)** the viral titers were measured via plaque assay. **(c)** mRNA fold change (log_2_) levels of several IFN response genes and **(d)** host cell-death related genes were determined by qRT-PCR from total RNA isolated 48 HPI from A549 cells infected with either H1N1 or H5N1 WT IAVs and treated with Combination (Baricitinib + Oseltamivir) 6 HPI. **(e)** A549 cells were infected with either H1N1 or H5N1 WT IAVs and treated with the combination either 6 or 12 HPI, total cell death was quantified via flow cytometry. Data was normalized with respect to No Drug control. Data are presented as mean ± SD from triplicate samples of a single experiment and are representative of results from three independent experiments. ***, p < 0.001, **, p < 0.01, and *, p < 0.05, by Brown-Forsythe and Welch ANOVA testes. Holm-Sidak’s multiple comparisons test (a and b) and two-way ANOVA with Sidak’s multiple comparison test (e).

### Delayed administration of Baricitinib and Oseltamivir combination restricts H1N1 IAV replication and associated pathology in the murine model

Encouraged by the initial results *in vitro*, we extended the study to evaluate the effectiveness of the drug combination against IAV infection in the C57BL/6 murine model. A dose of 10mg/kg body weight of each drug individually as well as in combination was found to be non-toxic (Supplementary Figure 6a). Under typical circumstances, Influenza symptoms usually appear after the infection is already well established, prompting patients to seek medical attention at that stage, or due to delayed diagnosis of the illness [39]. As demonstrated earlier, the combination treatment remains effective even when administered at later time point. Thus, we initiated the treatment in the murine model after the infection had occurred. Mice were infected intranasally with an LD_50_ dose of H1N1 IAV, and treatments were administered orally 1 or 2 days post-infection (DPI) (Figure 7a), and lung samples were harvested at day 4 post-infection. Treatment groups included: (i) Baricitinib, (ii) Oseltamivir and (iii) Combination of Baricitinib + Oseltamivir. Monitoring body weight and clinical signs (Figure 7b & c) revealed that animals treated with Baricitinib alone 1 DPI experienced a significant and early drop in body weight, falling below that of the infected control. This could have been due to early suppression of IFN signaling, impairing the innate immune response. However, when administered 2 DPI, Baricitinib alone was able to portray relatively lesser body weight drop as well as improved clinical outcomes. Oseltamivir administered alone at 1 DPI yielded the best results, consistent with its known antiviral activity thus effectively preventing body weight loss and improving clinical signs. However, when administered later at 2 DPI, a sharp decline in body weight was observed at 3 DPI that recovered only on 6 DPI, indicating reduced efficacy when treatment is delayed. Interestingly, the combination therapy was able to rescue the body weight and clinical signs when administered either 1 or 2 DPI. Furthermore, the lung viral titers measured at 4 DPI showed a 3-log reduction in animals treated with combination therapy at 2 DPI relative to infected control (Figure 7d). Combination therapy also achieved a 100% survival when administered either at 1 or 2 DPI (Figure 7e). Moreover, histopathological examination of infected lungs showed a significant reduction in pathology in the combination-treated group, as shown by the cumulative score (Figure 7f and g). Lungs from groups treated with the Combination showed less than 1. Perivascular infiltration 2. Perialveolar infiltration 3. Alveolar pneumonia and 4. Hemorrhage.

**Figure 7.**
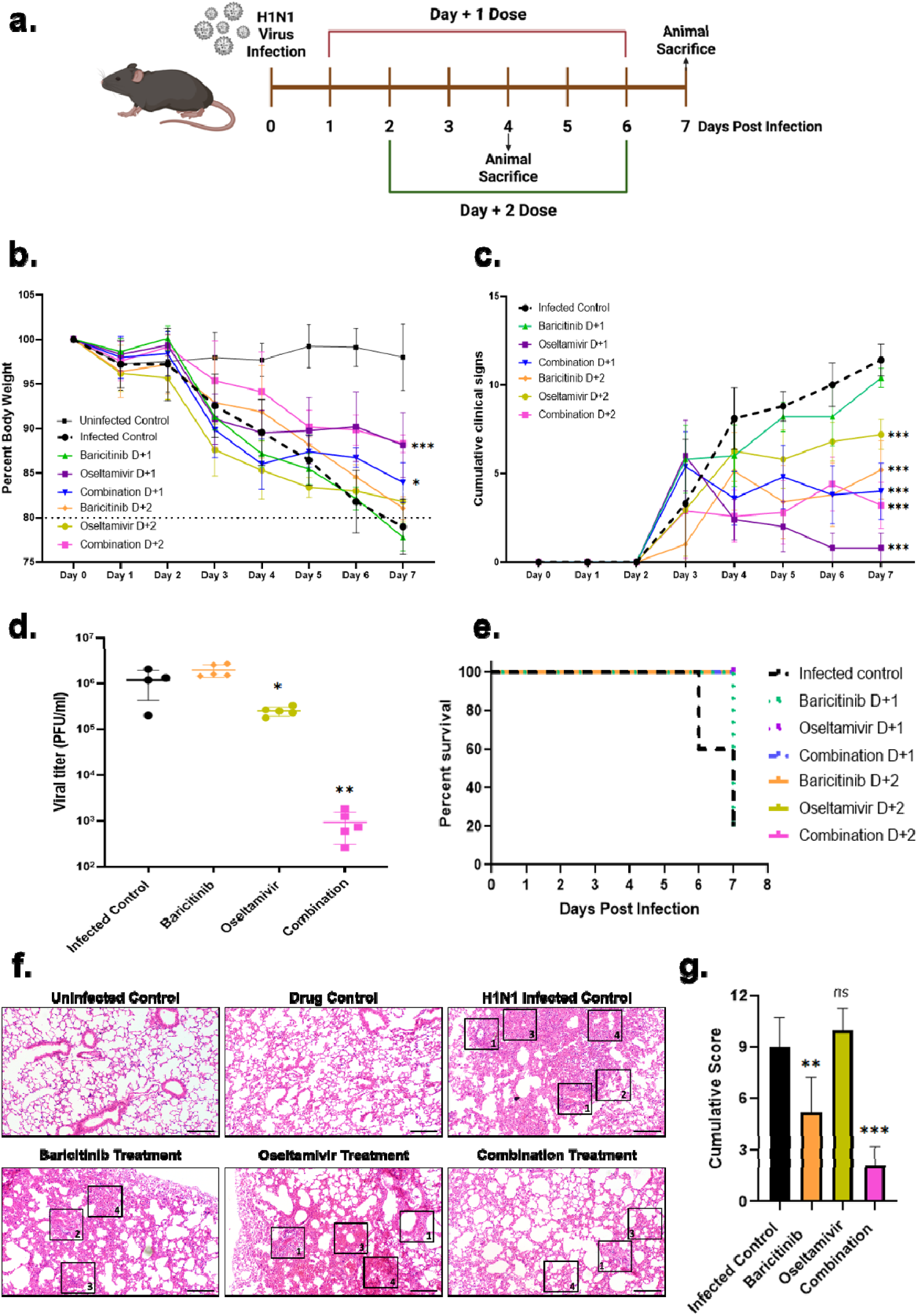
Delayed Combination therapy with Baricitinib and Oseltamivir mitigates H1N1 IAV replication and pathogenesis in murine model. **(a)** Schematic of drug treatment regime when mice were infected at LD_50_ of H1N1 virus and drug treatment started either D+1 or D+2 (10mg/kg body weight, in 0.5% CMC, once daily via oral gavage) and continued till D+7. (**b**) Percent body weight of the animals (**c**) Clinical signs [(i) Lethargy, (ii) Hunch-back, (iii) Piloerection and (iv) Rapid/abdominal breathing]. (**d**) Viral titers in lungs of animal treated with either of the drugs or their combination at 4 DPI (**e**) Survival of animals when treated with the drugs till 7 DPI. (**f and g**) Histology images for different groups and cumulative histology scoring (n = 5 per group). Criteria used for scoring and marked in the images include 1. Perivascular infiltration 2. Perialveolar infiltration 3. Alveolar pneumonia and 4. Hemorrhage. An objective histopathological scoring system was performed by a veterinarian blinded to study groups. Scale bar, 200 µm. Data are presented as mean ± SD from n=5 animals per group ***, p < 0.001, **, p < 0.01, and *, p < 0.05, by two-way ANOVA with Dunnett’s multiple comparisons test (b and c), One-way ANOVA with Dunnett’s multiple comparisons test (d) and one-way ANOVA with Holm-Sidak’s multiple comparisons test (g).

### Baricitinib and Oseltamivir combination also effectively reduces H5N1 IAV Replication and pathogenesis in a Murine Model

Based on these findings, we selected a 2 DPI treatment regimen to test against H5N1 IAV infection. Mice were infected intranasally with an LD_50_ of H5N1 IAV and started the treatment at day 2 DPI (Figure 8a). A sharp decline in body weight was observed around day 2 post-infection, consistent with earlier reports [17]. Analysis of body weight and clinical signs confirmed that the combination therapy provided superior protection when started 2 DPI (Figure 8b and c). Viral titers from lung samples harvested at 4 DPI showed an approximately 1-log reduction in the case of animals receiving combination therapy at 2 DPI compared to infected controls (Figure 8d). Combination-treated animals showed a 90% survival, outperforming other treatment groups (Figure 8e). Histopathological examination of infected lungs showed a reduction in pathology in the combination-treated group, as shown by the cumulative scores (Figure 8f and g). Lungs from groups treated with the Combination showed less than 1. Perivascular infiltration 2. Perialveolar infiltration 3. Alveolar pneumonia with Edema and 4. Hemorrhage. However, the improvements were less pronounced than in H1N1 IAV-infected mice. Excessive alveolar pneumonia with Edema was observed in the case of H5N1-infected lungs treated with either Baricitinib or Oseltamivir alone, but was somewhat alleviated by the combination treatment. The lung-to-body weight ratio was significantly reduced in animals treated with the combination therapy, suggesting a reduction in pulmonary edema (Supplementary Figure 7a). Lastly, the transcript levels of several cytokines were observed to be reduced when animals were treated with combination therapy (Supplementary Figure 7b). The therapeutic effects observed in animals infected with H5N1 were not as pronounced as those seen with H1N1, likely due to the markedly rapid replication rate of H5N1. This is evident from the sharp drop in body weight observed as early as 2 DPI in contrast to the 5-6 DPI decline seen in H1N1-infected animals. This suggests that the host system may be overwhelmed by the high viral burden at an early stage, limiting the ability of the treatment to effectively control the disease progression. It is possible that initiating the treatment earlier at 1 DPI may have yielded improved outcomes.

**Figure 8.**
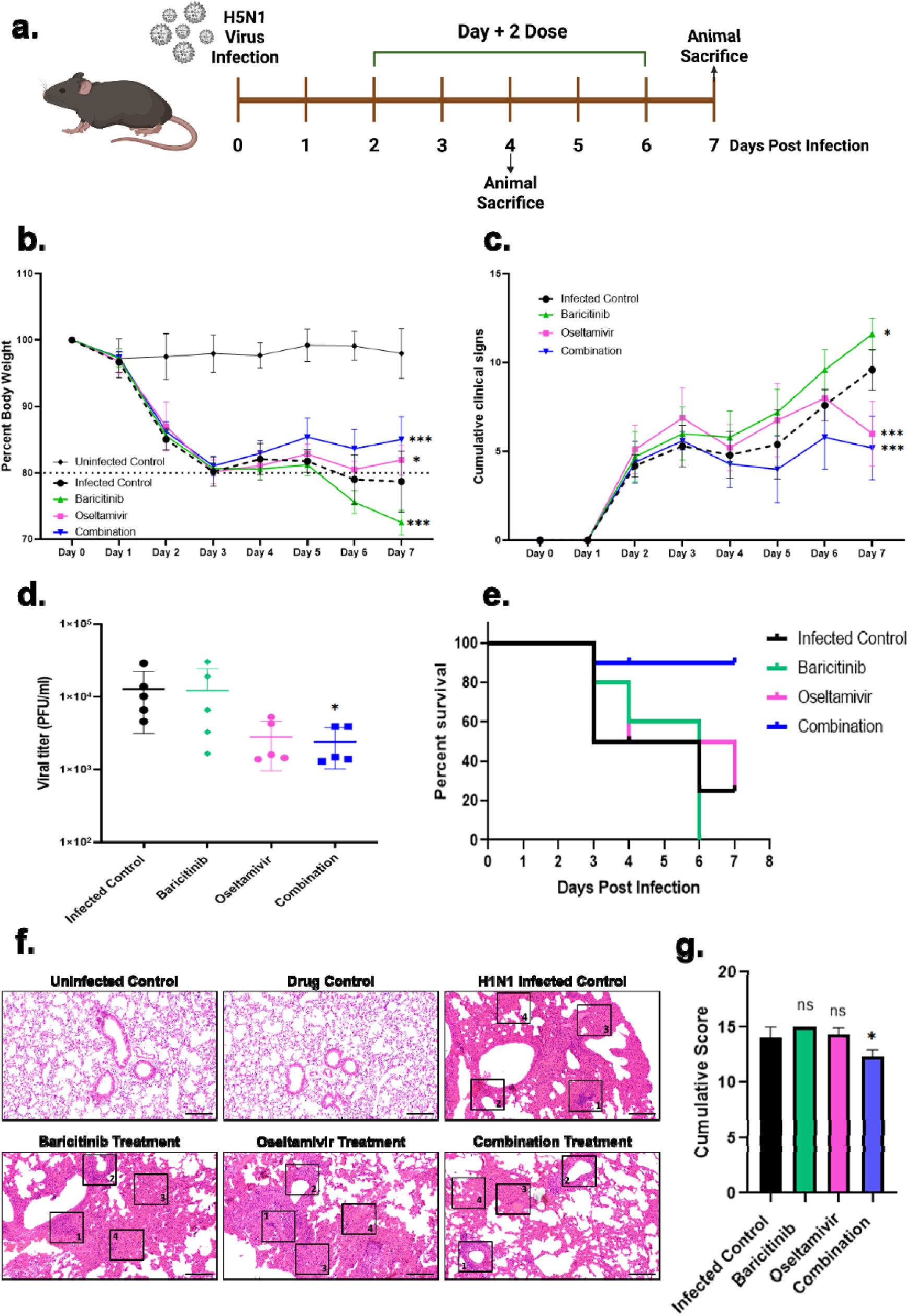
Delayed Combination therapy with Baricitinib and Oseltamivir effectively mitigates H5N1 IAV replication and pathogenesis in murine model. **(a)** Schematic of drug treatment regime when mice were infected at LD_50_ of H5N1 virus and drug treatment started D+2 (10mg/kg body weight, in 0.5% CMC, once daily via oral gavage) and continued till D+7. (**b**) Percent body weight of the animals **(c)** Clinical signs [(i) Lethargy, (ii) Hunch-back, (iii) Piloerection and (iv) Rapid/abdominal breathing] (**d**) Viral titers in lungs of animal treated with either of the drugs or their combination at 4 DPI (**e**) Survival of animals when treated with the drugs till 7 DPI. **(f and g)** Histology images for different groups and cumulative histology scoring (n = 5 per group). Criteria used for scoring and marked in the images include 1. Perivascular infiltration 2. Perialveolar infiltration 3. Alveolar pneumonia with Edema and 4. Hemorrhage. An objective histopathological scoring system was performed by a veterinarian blinded to study groups. Scale bar 200 µm. Data are presented as mean ± SD from n=5 animals per group ***, p < 0.001, **, p < 0.01, and *, p < 0.05, by two-way ANOVA with Dunnett’s multiple comparisons test (b and c), One-way ANOVA with Dunnett’s multiple comparisons test (d), one-way ANOVA with Holm-Sidak’s multiple comparisons test (g).

Furthermore, FluOMICS data [40] from C57BL/6 mice infected with either H1N1 or H5N1, with lung transcriptomics collected at 12 hours post-infection (HPI) and days 1, 2, 3, and 4 post-infection (DPI), were re-analyzed. The gene cluster map revealed distinct temporal gene expression patterns (Supplementary Figure 8a). Metascape analysis provided further insights where the inflammatory pathway was consistently enriched for H5N1-infected mice lungs, while the IFN-β peaked for H5N1 around 1 DPI, and it came down by 4 DPI (Supplementary Figure 8b). Within this dataset, inflammatory cytokines and cell death were specifically curated and compared (Supplementary Figure 8c). It was observed that most cytokines peaked at 2 DPI, suggesting this as a logical time point to initiate anti-inflammatory treatment. Additionally, IFN-β expression peaked around 1 DPI in H5N1-infected mouse lungs but subsequently declined, becoming lower than that in H1N1-infected lungs by 4 DPI. Overall, our murine experiments suggest that neither Baricitinib nor Oseltamivir alone effectively controls IAV infection and associated inflammation in H1N1 or H5N1 infections. However, their combination significantly reduces viral load, mitigates lung inflammation, improves survival, and alleviates clinical symptoms. Thus, these findings suggest that this combination therapy holds a strong clinical potential for managing influenza during epidemics or pandemics.

## Materials & Methods

### Ethics Statement

This study was conducted in compliance with institutional biosafety guidelines (IBSC/IISc/ST/17/2023), following the Indian Council of Medical Research and Department of Biotechnology recommendations. All Influenza A virus animal work was performed in a Viral Biosafety Level-3 facility. The experiments were performed according to CPCSEA (The Committee for Control and Supervision of Experiments on Animals) guidelines.

### Cell Lines and Virus

A549 (NCCS, Pune, India) and MDCK (NCCS, Pune, India) cells were cultured in complete Dulbecco’s modified Eagle medium (12100-038, Gibco) with 10% HI-FBS (16140- 071, Gibco), 100 IU/mL Penicillin, and 100 mg/mL Streptomycin (15140122, Gibco) supplemented with GlutaMAX (35050-061, Gibco). Influenza A viruses A/California/04/2009 (H1N1) [H1N1] and A/Viet Nam/1203/2004 (H5N1) [H5N1] (with polybasic site eliminated - HALo) wild type, as well as GFP reporter viruses, were a kind gift from Prof. Adolfo Garcia-Sastre (ISMMS, NY). A/Northern Shoveler/Mississippi/11OS145/2011 (H7N9), A/Mallard/Wisconsin/2576/2009 (H5N1) were obtained from BEI resources. A/Wisconsin/15/2009 (H3N2) was obtained from ATCC. These viruses were propagated in 11-day-old embryonated chicken eggs or MDCK cells and titrated by plaque assay in MDCK cells as reported previously [41]. Briefly, MDCK cells were seeded in a 6-well plate and allowed to grow to full confluency. Viral stocks were serially diluted 10-fold in Opti-MEM (31985070, Gibco). Cells were washed twice with 1X PBS (20-031-CV, Corning) and the viral inoculum was added onto them. The plates were rocked from side to side every 10 mins, and after 1 hour, the inoculum was replaced with plaque assay overlay media (Plaque assay media + H_2_O + 1% DEAE Dextran (D9885, Sigma) + 1ug/ml TPCK trypsin (T1426, Sigma Aldrich) + 2% Oxoid agar (LP20028, Thermo Fischer Scientific)) where Plaque assay media contains 2X MEM (61100061, Gibco) + 0.3% BSA (160069, MP Biologicals) + Sodium bicarbonate (195494, MP Biologicals) + Pen-Strep + HEPES (H5303, Promega). 48 hours post infection, the cells were fixed with 4% formaldehyde (Q24005, Qualigens) followed by staining of the plaques (H_2_O + Coomassie Brilliant blue (B-0149, Sigma) + Methanol (Q34457, Qualigens)).

### Virus infection of cell culture

A549 cells in 24-well plates were infected with 100 µl per well with 1 or 0.1 MOI of H1N1 or H5N1, either Wild type (WT) or GFP reporter virus, in Opti-MEM in triplicate. The plates were rocked from side to side every 10 mins, and after 1 hour, the inoculum was replaced with infection medium (1X MEM containing 0.3% BSA and 0.1 µg/mL TPCK trypsin). Cells were harvested as per the experimental requirements.

### Fluorescence and Brightfield Microscopy

At the desired time point post infection, Propidium iodide (556463, BD) was added to the media directly at least 1 hour prior to subjecting the cells to fluorescent or brightfield microscopy using Evos M5000 (Thermo Fisher Scientific). All the images were analyzed using ImageJ Fiji.

### MTT Assay

Cell viability of uninfected vs infected cells was measured using the MTT (M2128, Sigma) assay as per the manufacturer’s instructions. Briefly, cells were seeded and infected in 96- well format. After the desired time point, cells were washed with 1X PBS, and 0.4 mg/ml MTT was added to each well. Cells were incubated at 37°C for 4 hours, followed by adding DMSO to each well and measuring the absorbance at 590 nm with a reference at 630 nm.

### PI Staining and Flow Cytometry

1 X 10^5^ adherent cells were stained with PI staining kit (560506, BD) as described previously [42]. Briefly, floating dead cells in the supernatant from the infected/treated wells were collected in a 15 ml tube. Remaining adherent cells were then washed with cold PBS and trypsinized (25200072, Gibco). The detached cells were then pooled with the floating cells, pelleted, and resuspended in 1X PBS. 1ul of PI each was added to the suspension and incubated for 15 mins at room temperature (RT) in the dark. 400 µl of PBS was added to each tube and analyzed via flow cytometry – CytoFlex (Beckman Coulter).

### Cell Cytotoxicity

Baricitinib (HY-15315, MedChemExpress) and JAK Inhibitor I (420099, Merck Millipore) were resuspended in DMSO (D2650-100ML, Sigma-Aldrich), while Oseltamivir (HY-17016, MedChemExpress) was resuspended in water. Cytotoxicity of the drugs Baricitinib, JAK Inhibitor I, and Oseltamivir was tested in A549 cells. Cells were seeded in a 96-well plate and were treated with increasing concentrations of the drugs ranging from 0-50 μM for 48 hours. Later, the cells were lysed using the Cell Titer Glo cell viability kit (Promega Cat# G7570), and the luminescence was measured using a Tecan Plate reader (INFINITE M PLEX).

### Drug treatment

A549 cells were infected with WT or GFP reporter H1N1 and H5N1 virus, 6 HPI cells were treated with 1 μM of either Baricitinib or JAK inhibitor I. 48 HPI cells were visualized under a fluorescent microscope or harvested either for flow cytometric analysis or western blotting. Oseltamivir was used in combination with Baricitinib for cells infected with different IAVs, 6HPI. In the combination experiments, 0.5 μM of Baricitinib and 1 μM of Oseltamivir were used.

### Luciferase Reporter Assay

To test the IFN-β promoter activity (i.e., IFN induction), HEK-293 T cells (0.1 X 10^6^ cells/well in a 24-well plate) were co-transfected, in duplicates, with 50 ng of pIFNβ-luc Firefly luciferase reporter plasmid and 20 ng of pRL-TK Renilla luciferase reporter plasmid (as internal control). For the ISRE promoter activity (i.e., signaling), cells were co-transfected with 50 ng of ISRE-luc Firefly Luciferase reporter plasmid and 20 ng of pRL-TK Renilla Luciferase reporter plasmid. 24 h post-transfection, cells were infected with 1 MOI H1N1- WT, H5N1-WT, H1N1-GFP, or H5N1-GFP IAVs. 16 hours post-infection, cells were lysed and luciferase activity was measured using a dual-luciferase assay system (Promega Cat# E1980) according to the manufacturer’s instructions. Firefly and Renilla Luciferase signals were quantified using Tecan plate reader (INFINITE M PLEX). The signals were represented as percentage induction with respect to uninfected cells.

### Generation and validation of STAT1-/- A549 cells

Using the CRISPR-Cas9 system, a targeting guide DNA was delivered into A549 cells by transfection of the pSp-Cas9-2A-GFP-gDNA plasmid. The GFP-positive cells were sorted using BD FACS Aria Fusion Sorter (Central FACS Facility, IISc, Bengaluru) and grown as single-cell colonies. Several colonies were screened by western blotting for STAT1 and by RT-PCR for a few Interferon Stimulated Genes (ISGs) upon interferon treatment (1000 Units/ml) to identify colonies truly knocked out of STAT1. Verified STAT1 -/- colonies were expanded and used for experiments.

### qRT-PCR

Cells were harvested in TRIzol (Thermo Fisher, 15596018) as per the manufacturer’s instructions. 1 mg of RNA was reverse transcribed into cDNA using Prime Script^TM^ RT Reagent Kit with gDNA Eraser (Perfect Real Time - RR047A, Takara-Bio) and then diluted 5-fold with nuclease-free water (112450204, MP Biomedicals). The gene expression study was conducted using PowerUp^TM^ SYBRTM Green Master Mix (A25742, Applied Biosystems^TM^) with 18S rRNA as the internal control and appropriate primers for the genes using Quant Studio 5 (Applied Biosystems, Thermo Fisher Scientific)

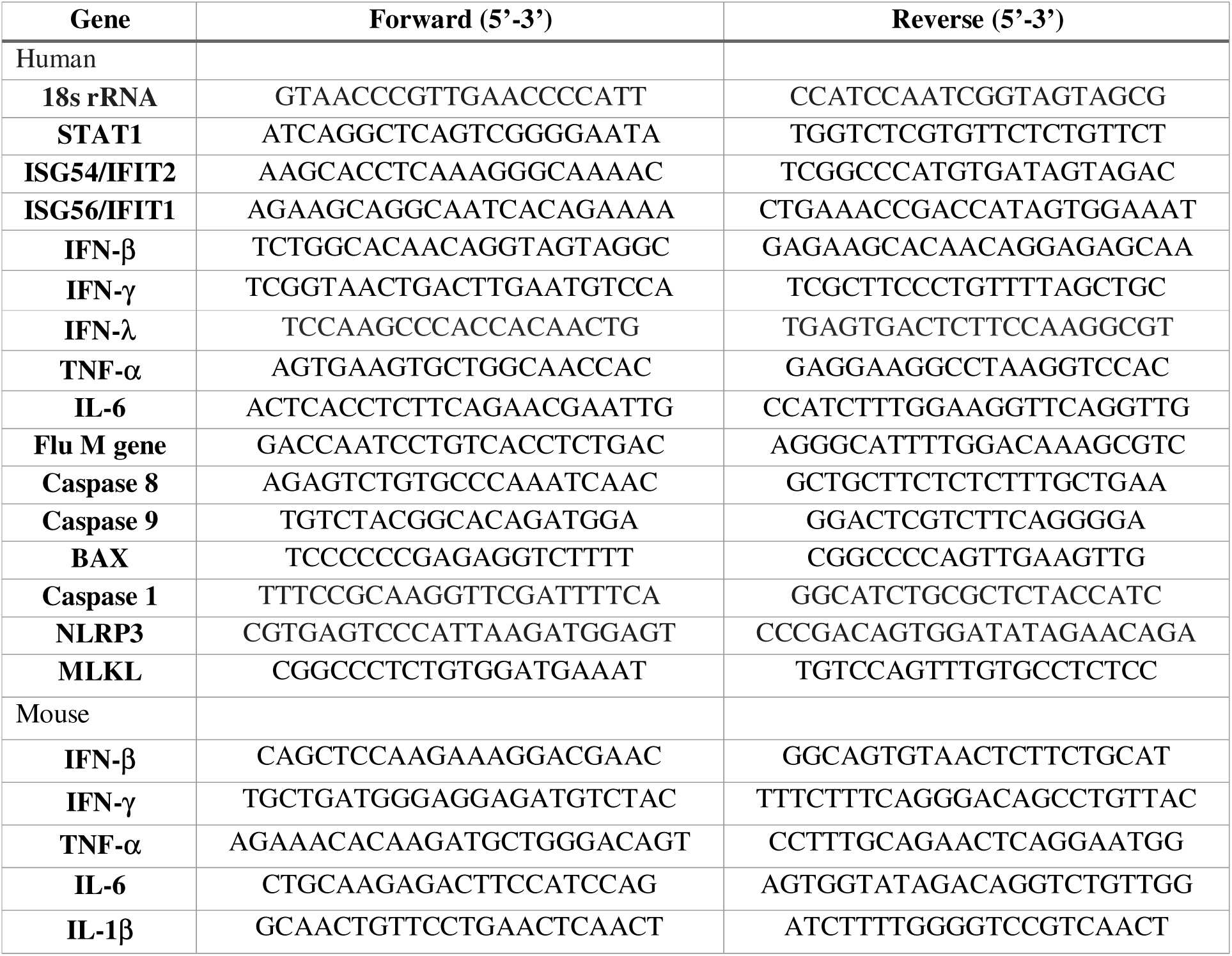

### Western blotting

Cells were washed with 1X PBS, lysed with 1x Laemmli buffer (1610747, BIO-RAD), and heated at 95°C for 10 min. Cell lysates were then subjected to standard SDS-PAGE, and separated proteins were transferred onto a PVDF membrane (IPVH00010, Immobilon-P; Merck). The membrane was incubated in a blocking buffer containing 5% Skimmed milk (Sigma Aldrich, 70166) in TBS containing 0.05% Tween 20 (Sigma-Aldrich P1379) (1X TBST) for 2 h with slow rocking at room temperature (RT). Primary antibodies, Pan Influenza NP (ab128193, Abcam), Caspase 3 and cleaved caspase 3 (NB100-56708, Novus Biologicals) Caspase 8 and cleaved Caspase 8 (4790S, CST), Gasdermin-D and cleaved Gasdermin-D (97558T, CST) and β actin (Sigma Aldrich, A1978) incubation in BSA was done for 12-14 hours at 4°C with gentle rocking, after which the membrane was washed with 1X TBST and incubated for 2 h with secondary antibody, goat-anti rabbit (Thermo Fisher Scientific, 31460) and goat-anti mouse (Thermo fisher Scientific, 31430) in blocking buffer at RT. After a further wash with 1X TBST, the blots were developed using Clarity Western ECL Substrate (Bio-Rad, 1705061).

### Bioinformatics

#### (i) Differential Gene Expression Analysis

Transcriptomics data were obtained from the FluOMICS data resource “RNA-Seq transcriptome profiling of human tracheobronchial epithelial (HTBE) cells in response to infection with Influenza A/California/04/09 (H1N1), A/Wyoming/03/03 (H3N2), and A/Vietnam/1203/04 (H5N1) H5N1.” [34], or of C57BL/6 mice lungs in response to infection with the above-mentioned IAVs [40]. Samples were collected at different time points: 3, 6, 12, and 18 hours for HTBE cells with their respective control at each point. In the case of mouse lung tissue, samples were similarly harvested from three different mice at the time points of 12 hours, 1 day, 2 days, 3 days, and 4 days. Data for H1N1 and H5N1 were used for further analysis. Metadata was downloaded for all samples using the corresponding accession numbers. Sample identifiers were matched with experimental conditions (infected vs. mock, time points, and virus strain). Quality control metrics were assessed to ensure data integrity before proceeding to downstream analysis. Raw gene count files were acquired for each sample to quantify gene expression levels. The BioJupies platform was used to download the count files. Each file contained raw counts representing the number of reads mapping to each gene for each sample. Data was organized into a matrix for subsequent analysis in R. Raw count files were imported into the DESeq2 pipeline in R. Low-expressed genes were filtered out to reduce noise. Counts were normalized to account for differences in sequencing depth and RNA composition across samples. Differential expression analysis was performed using DESeq2 to compare infected samples against their respective mock controls at each time point. Genes were considered significantly differentially expressed if they met the following criteria: Log_2_ Fold Change (LFC) ≥ 3.32 and p-Value ≤ 0.01. These genes are considered to have significant and biologically meaningful changes in expression in response to viral infections. A cluster map was plotted as a heat map using the Python Seaborn library. Using this as a reference, the filtered data was curated for the presence of these genes and further plotted as a heat map as a function of their differential expression using the R package ggplot. A cytokine-related gene list was obtained by querying the term “cytokines” in the GeneCards database. This list was filtered with the set of upregulated genes in the final long-format dataset to generate a focused heatmap of cytokine gene expression. For the mouse-specific analysis, cytokine genes were identified using the Mouse Genome Informatics (MGI) database. While pathway analysis was plotted as a heat map of the top 20 enriched terms, by Metascape [43] (https://metascape.org/gp/index.html#/main/step1).

### Animal experiments

Mice were used for the experiments because they are the most widely used preclinical animal models for IAV research [44]. For IAV infection experiments, 6-8 weeks old, female C57BL/6 mice (Central Animal Facility, Indian Institute of Science, Bengaluru, India) were used. All animals were housed in groups of five in individually ventilated cages maintained at 23 ± 1°C temperature and 50 ± 5% relative humidity. The animals were given standard pellet feed and water *ad libitum* and maintained on a 12-h day/night light cycle at the Viral Biosafety Level-3 facility, Indian Institute of Science. All animals were monitored daily during the experiment. An overdose of Ketamine (Bharat Parenterals Limited) and Xylazine (Indian Immunologicals Ltd) was used to sacrifice animals upon completion of the experiment.

### Drug treatment and infection in a murine model

The drugs were resuspended in 0.5% Carboxy methyl cellulose (CMC) (C4888-500G, Sigma). CMC was added to lukewarm water followed by mixing at 37°C. Literature suggested the use of 10mg/kg body weight of Baricitinib [45, 46] and Oseltamivir [47, 48] individually which corresponds to a Human Equivalent Dose (HED) of 10mg/kg body weight (Mouse) * 0.08 (Conversion factor) = 0.8mg/kg each. Conversion is as per FDA guidelines https://www.fda.gov/media/72309/download. To assess the toxicity of the combination, animals were treated orally with either Baricitinib, Oseltamivir, or their combination for 12 days. The *in vivo* half-life of Baricitinib is 12.5 hours; hence, it was given once daily [49]. However, Oseltamivir is often given twice daily [47, 48] which has been reduced to once daily in this study. The body weight and general health of the animals were measured every day for up to 12 days post-treatment.

Infection groups were divided into 4: (i) Infected control, (ii) Baricitinib, (iii) Oseltamivir, and (iv) Combination (Baricitinib + Oseltamivir). For infection, mice under IP anesthesia with Ketamine (90 mg/kg) and Xylazine (4.5 mg/kg) were challenged intranasally with the LD_50_ of either H1N1 or H5N1 IAV virus in 40 μL PBS. They were treated with the drugs at Day +1 or 2 post-infection. Clinical signs including (i) Lethargy, (ii) Hunch-back, (iii) Piloerection, and (iv) Rapid/abdominal breathing were considered. Scoring was done based on the severity on a scale of 1–3 (1-mild, 2-moderate, 3-severe). Half of the animals were sacrificed 4 days post infection and lungs were collected for plaque assay, qRT-PCR, and histology, remaining were sacrificed 7 DPI. For plaque assay, lung samples were collected in MEM containing 0.3% bovine serum albumin (Sigma, A7906), homogenized, and centrifuged at 5000xg for 10 min at 4°C to pellet tissue debris. The supernatant was used for plaque assay as described previously. For qRT-PCR, lung samples were collected in Trizol and homogenized followed by qRT-PCR as described previously.

### Histopathology

Mice lung tissue samples were fixed in 10% buffered paraformaldehyde (PFA), embedded in paraffin blocks, and tissue sections of 4-6 µm thickness were made using a microtome. The sections were then stained with Hematoxylin and Eosin and examined by light microscopy as previously described [50]. Clinical scoring for infected mice lung was done using 4 different pathological criteria: (i) Perivascular infiltration, (ii) Perialveolar infiltration, (iii) Alveolar pneumonia & Edema, and (iv) Hemorrhage. Scoring was done based on the severity on a scale of 1–3 (1-mild, 2-moderate, 3-severe). Histopathology analysis and scoring were done by a trained veterinary pathologist who was blinded to treatment groups.

### Quantification and statistical analysis

All statistical analyses were performed using GraphPad Prism 8.4.3 (GraphPad Software, USA). Details about the statistical method used are mentioned in the legend section of the respective figures. Error bars indicate mean ± SD from triplicate samples. In all cases, a p-value <0.05 is considered significant.

## Discussion

Influenza virus infection remains a significant global health challenge. It contributes not only to substantial mortality but also to severe morbidity, leading to a considerable economic burden annually [51]. Strategies to combat influenza primarily fall into two categories: prevention and treatment. The mainstay of prevention is vaccination; however, its effectiveness is limited by the virus’s high mutation rate, particularly in the hemagglutinin (HA) protein. This necessitates continual prediction and reformulation of vaccines every 6– 12 months, often with suboptimal efficacy [52]. The development of a universal influenza vaccine remains an unmet goal [53].

On the therapeutic front, antiviral agents, particularly neuraminidase inhibitors such as oseltamivir and zanamivir, are widely used. However, the influenza virus can rapidly develop resistance due to its high mutation rate, particularly in the regions targeted by these drugs [54, 55]. Consequently, there is a pressing need to develop alternative therapeutic strategies that can effectively reduce disease burden and are less susceptible to viral escape. One promising approach is to target host pathways rather than viral components, which may be less prone to resistance due to their host origin. Of the various combinations of hemagglutinin (H1–H18) and neuraminidase (N1–N11), only H1N1 and H3N2 subtypes are currently endemic in humans. Sporadic zoonotic transmissions from avian viruses such as H5N1 and H7N9 have been reported [56]. The 2009 H1N1 pandemic virus (H1N1pdm09) was highly transmissible but associated with a relatively low case fatality rate. In contrast, avian H5N1 infections, although rare in humans and not efficiently transmitted from person to person, exhibit high fatality rates and pose a potential pandemic threat [57] [58]

Severe H5N1 infection is characterized by extensive lung damage, largely attributed to excessive cell death, vascular leakage, edema, and tissue destruction [59]. In our study, we investigated virus-induced cell death and host responses in cells infected with H1N1 and H5N1 viruses. We observed that H5N1 replicated more rapidly and induced significantly higher levels of interferon-beta (IFN-β), along with accelerated and extensive cell death.

Although type I interferons (IFN-α/β) play a critical role in antiviral defense, their uncontrolled production can trigger a harmful cytokine storm, contributing more to tissue damage than to viral clearance [60]. In pediatric influenza cases, severe outcomes are often associated with an excessive cytokine response rather than high viral load. This may explain the limited efficacy of antiviral drugs like neuraminidase inhibitors in children [61]. Thus, a primary therapeutic goal in such cases is to modulate the immune response, particularly by controlling cytokine storm–induced inflammation.

Typically, higher IFN-β levels correlate with lower viral replication due to enhanced antiviral states. However, prior studies have demonstrated that the role of IFN-α/β in antiviral gene induction within epithelial cells may be redundant, with STAT1 playing a more pivotal role. Furthermore, IFN-α/β has been shown to drive immunopathology in susceptible hosts, independent of viral load [62]. Our findings show, that H5N1 infection resulted in elevated IFN levels without a corresponding reduction in viral titers. This suggests that uncontrolled viral replication may drive excessive interferon production, which in turn fails to contain the virus and exacerbates inflammation and tissue injury. Interestingly, H1N1-infected cells exhibited a different profile—reduced yet sustained IFN production, accompanied by significant bystander cell death. This pattern may limit viral spread to adjacent cells while minimizing inflammation. Moreover, H1N1 predominantly induced apoptotic (non-inflammatory) cell death, whereas H5N1 triggered more inflammatory forms of cell death, likely contributing to lung pathology. Given the prominent role of IFN signaling in influenza-induced cell death, we employed STAT1 knockout (KO) cells to assess its contribution. STAT1 deficiency led to a marked reduction in cell death for both viruses, with a more pronounced effect observed in H1N1-infected cells. This is likely due to the lower baseline IFN production during H1N1 infection and the further attenuation of signaling in the absence of STAT1. Similar results were obtained using Baricitinib, a JAK inhibitor and FDA-approved anti-inflammatory drug for rheumatoid arthritis, which also blocks IFN signaling.

This protective effect of Baricitinib was observed across multiple strains, including human H1N1, H3N2, avian H5N1, and H7N9 viruses, suggesting a broadly applicable mechanism. However, STAT1 inhibition alone resulted in increased viral replication, indicating that IFN blockade, while reducing inflammation, compromises antiviral control. Thus, we explored the combined use of Baricitinib with the direct-acting antiviral Oseltamivir. *In vitro* experiments demonstrated that the combination of Baricitinib and Oseltamivir significantly reduced both viral titers and cell death. Encouraged by these results, we extended the study to a mouse model. In combination therapy, we administered oseltamivir once daily—half the typical dosage—and paired it with Baricitinib [47]. Our *in vitro* preliminary data also suggest that Baricitinib concentrations can be reduced without compromising efficacy, though this remains to be validated *in vivo*.

To mimic clinically relevant conditions, we initiated treatment two days post-infection. This delayed regimen was effective in improving survival in H1N1-infected mice, achieving 100% survival, and showed 90% survival in H5N1-infected mice—surpassing survival rates reported in previous combination therapy studies for H1N1pdm (∼87%) [63], for H1N1 (PR/8) (80%) [64] and for H5N1 (53.3%) [65]. Although IFN levels declined by day 4 post-infection in H5N1-infected mice, peak production and associated weight loss occurred earlier (around day 2), consistent with the transient yet intense cytokine surge seen in H5N1 infections. Despite lower IFN levels at later stages, early IFN overproduction was sufficient to cause extensive lung damage, as confirmed by histopathological analysis. Our findings underscore the pathogenic role of dysregulated IFN-β production in severe influenza infections, particularly H5N1, and highlight the therapeutic potential of combining host-directed anti-inflammatory strategies with direct-acting antivirals. While IFN signaling blockade alone is insufficient due to increased viral replication, its combination with oseltamivir shows promise in both limiting viral load and mitigating tissue damage.

Importantly, since both drugs are FDA-approved, their repurposing for influenza treatment offers a feasible path toward clinical application. While previous studies have explored combination therapies for influenza, additional approaches remain valuable to broaden therapeutic options. Blocking IFN signaling can have a wide range of effects, with the regulation of cell death being just one aspect. In this *in vivo* study, we provide indirect evidence that IFN signaling inhibition may contribute to reduced cell death and, consequently, decreased inflammation. However, this observation warrants further investigation to elucidate the underlying mechanisms and broader implications. Nonetheless, this strategy represents a promising step toward more effective and adaptable influenza therapies.

## Acknowledgments

ST acknowledges funding support from DBT-Wellcome Trust India Alliance (IA/I/18/1/503613) and ICMR (IIRPIG-2023-0000978). ST acknowledges infrastructure funding from DST-FIST, BIRAC, BFI, and IISc. MS and RTY acknowledge the MOE fellowship. RN is supported by the DBT RA fellowship. SS acknowledges the PMRF fellowship.

## Author Contribution

Conceptualization, Funding acquisition, Project administration, Supervision, Resources: AGS, ST

Methodology, Data curation, Formal Analysis, Validation, Visualization: MS, AK, DB, RTY, RN, SS, GW

Manuscript Writing – review & editing: MS, ST

## Declarations

### Consent for publication

All the authors have provided consent for publication.

### Data Availability

All primary data associated with this study have been included in the manuscript. Any additional information queries can be directed to the corresponding author.

---

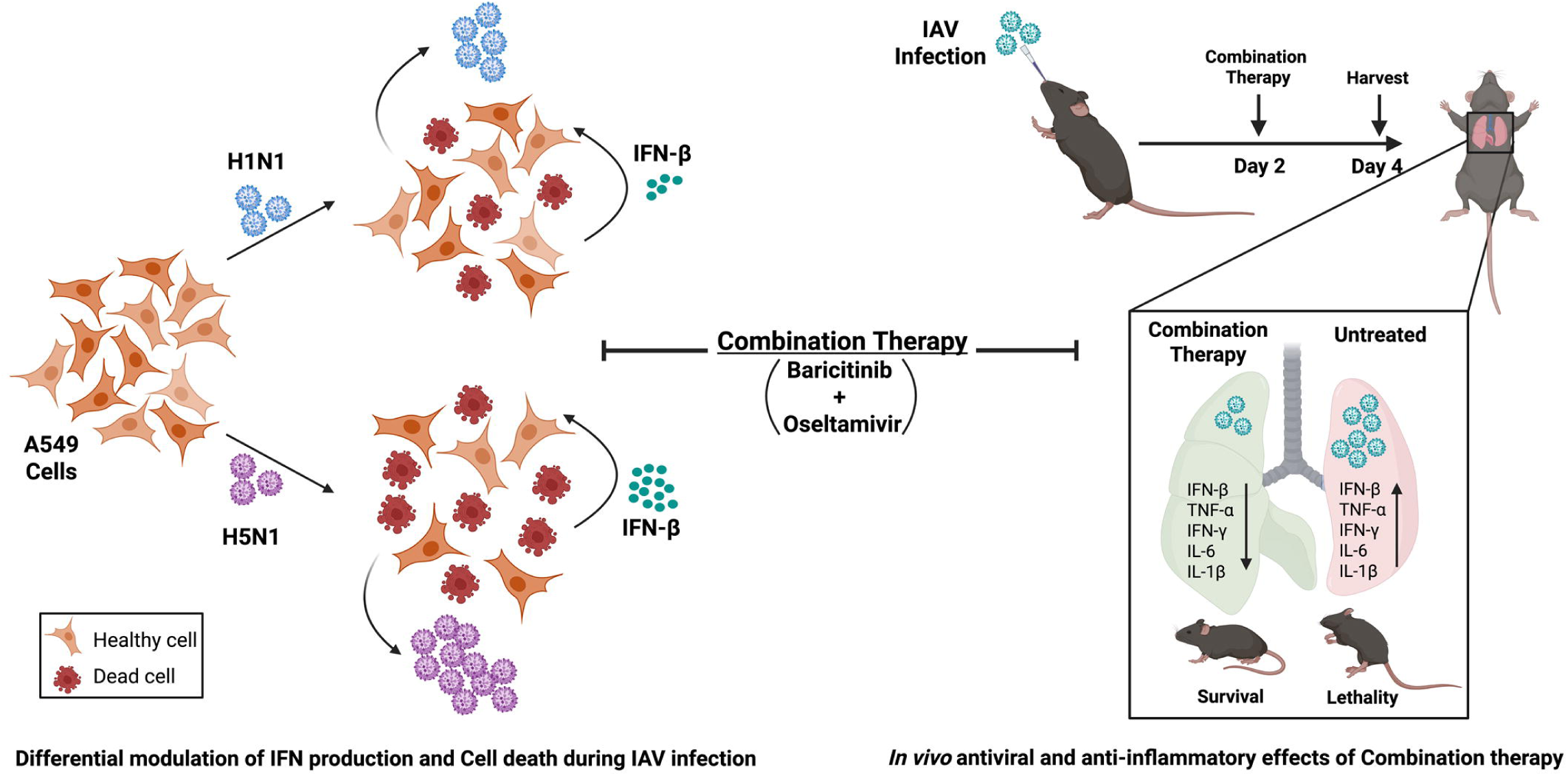

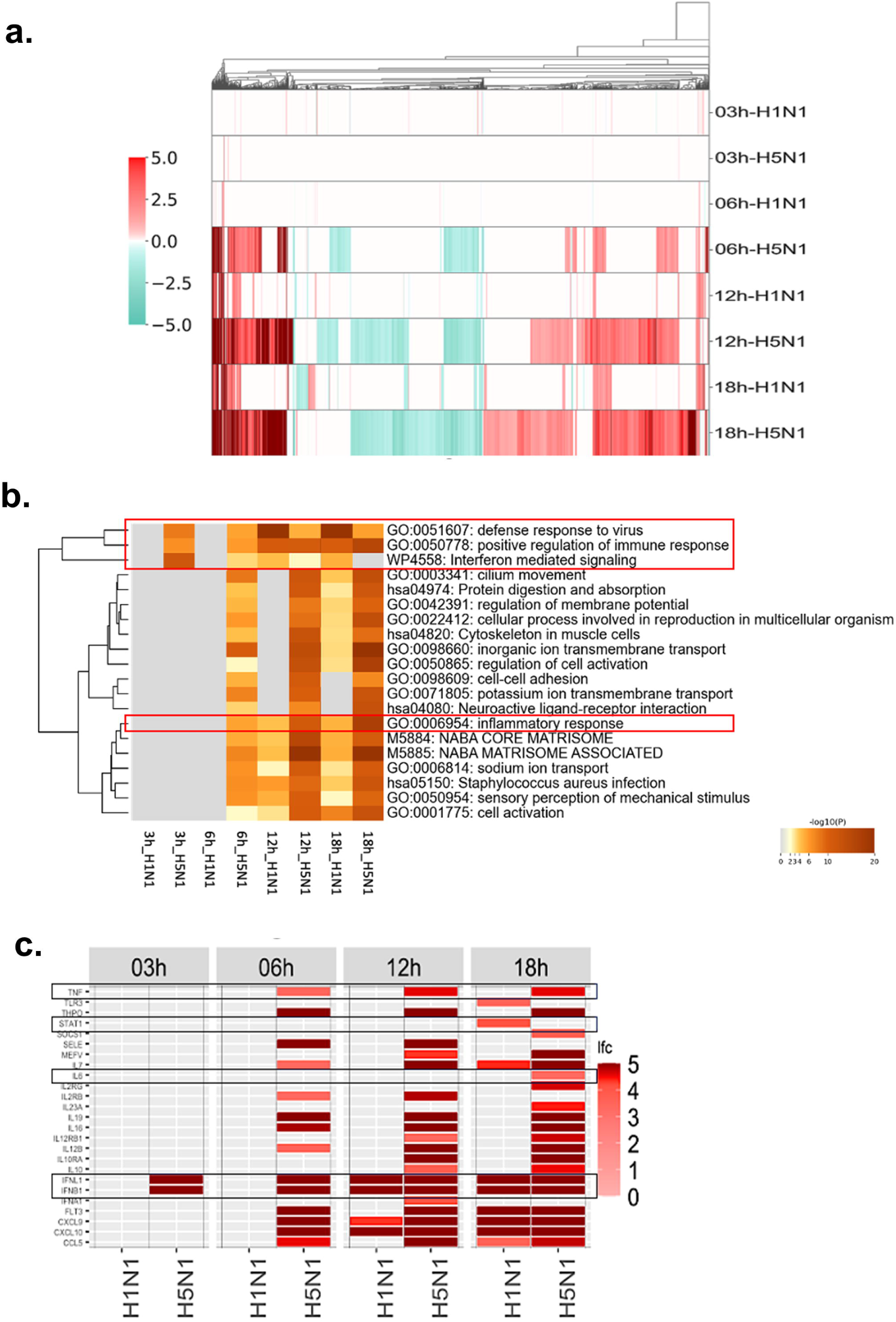

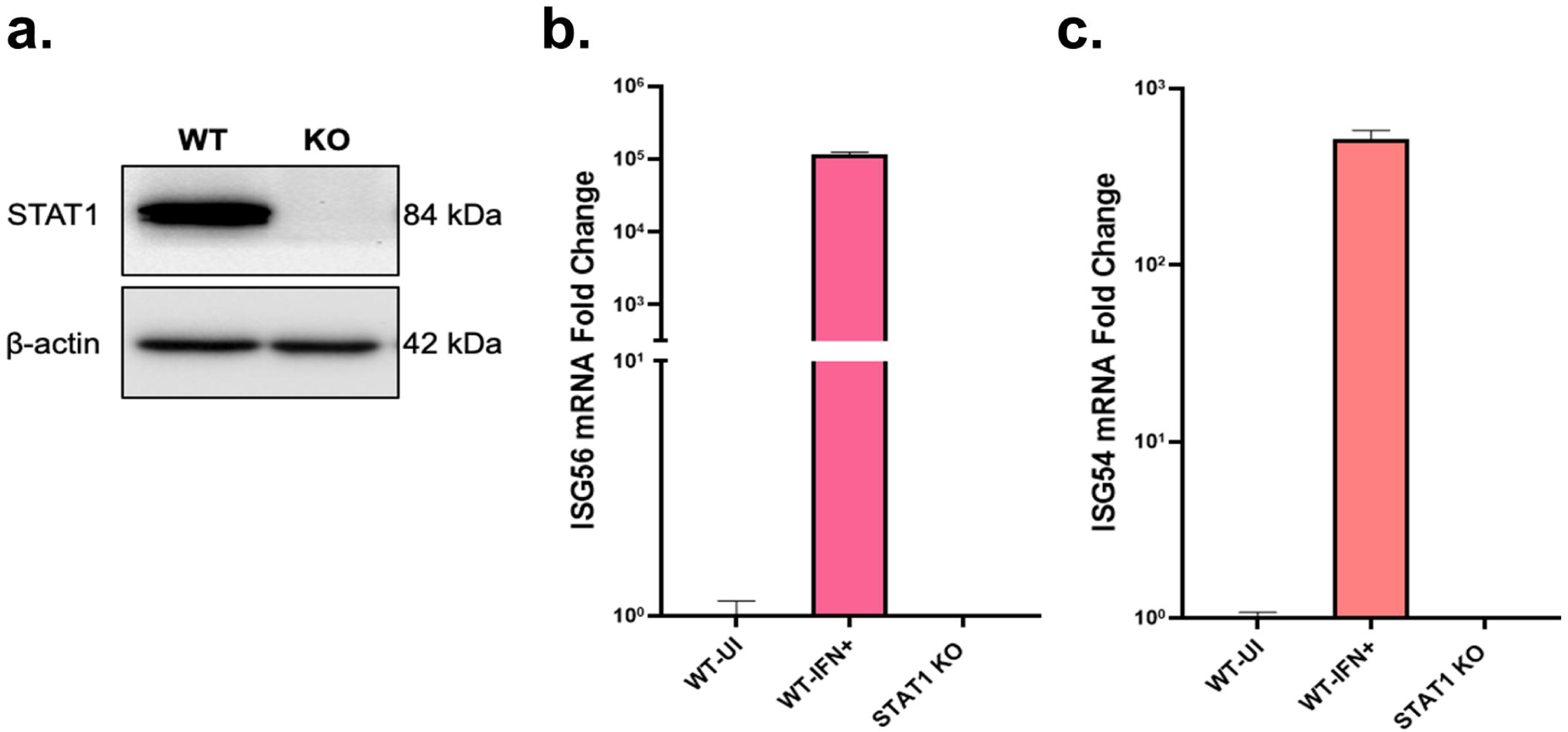

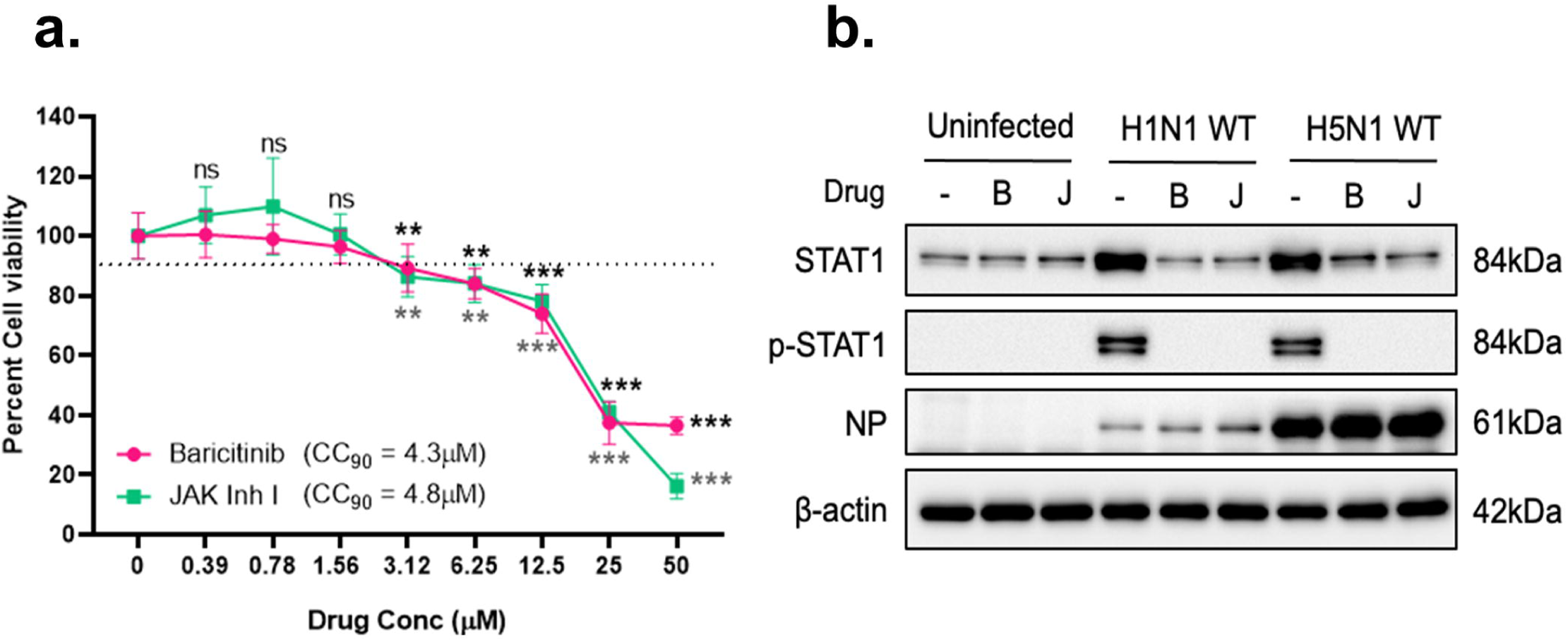

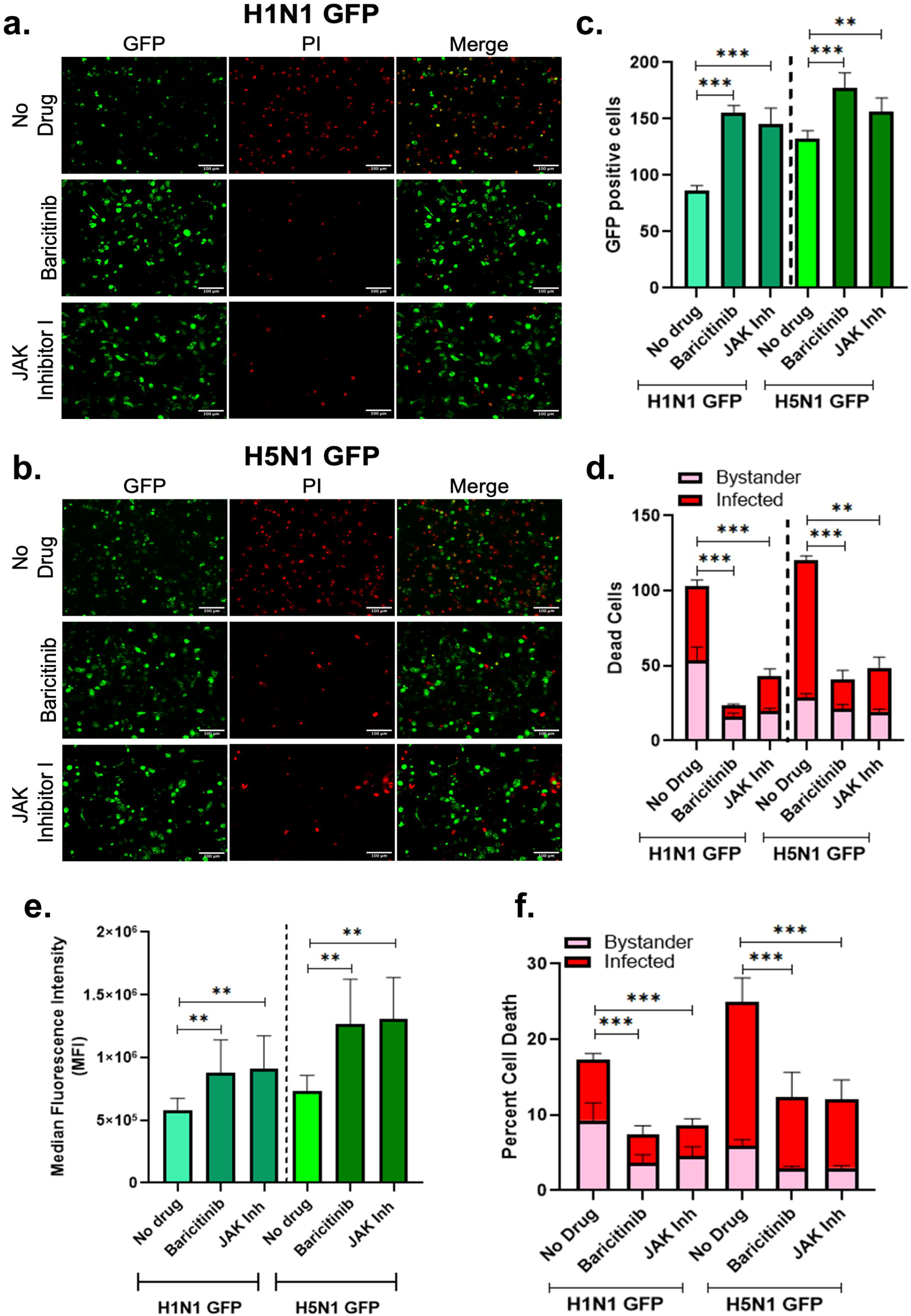

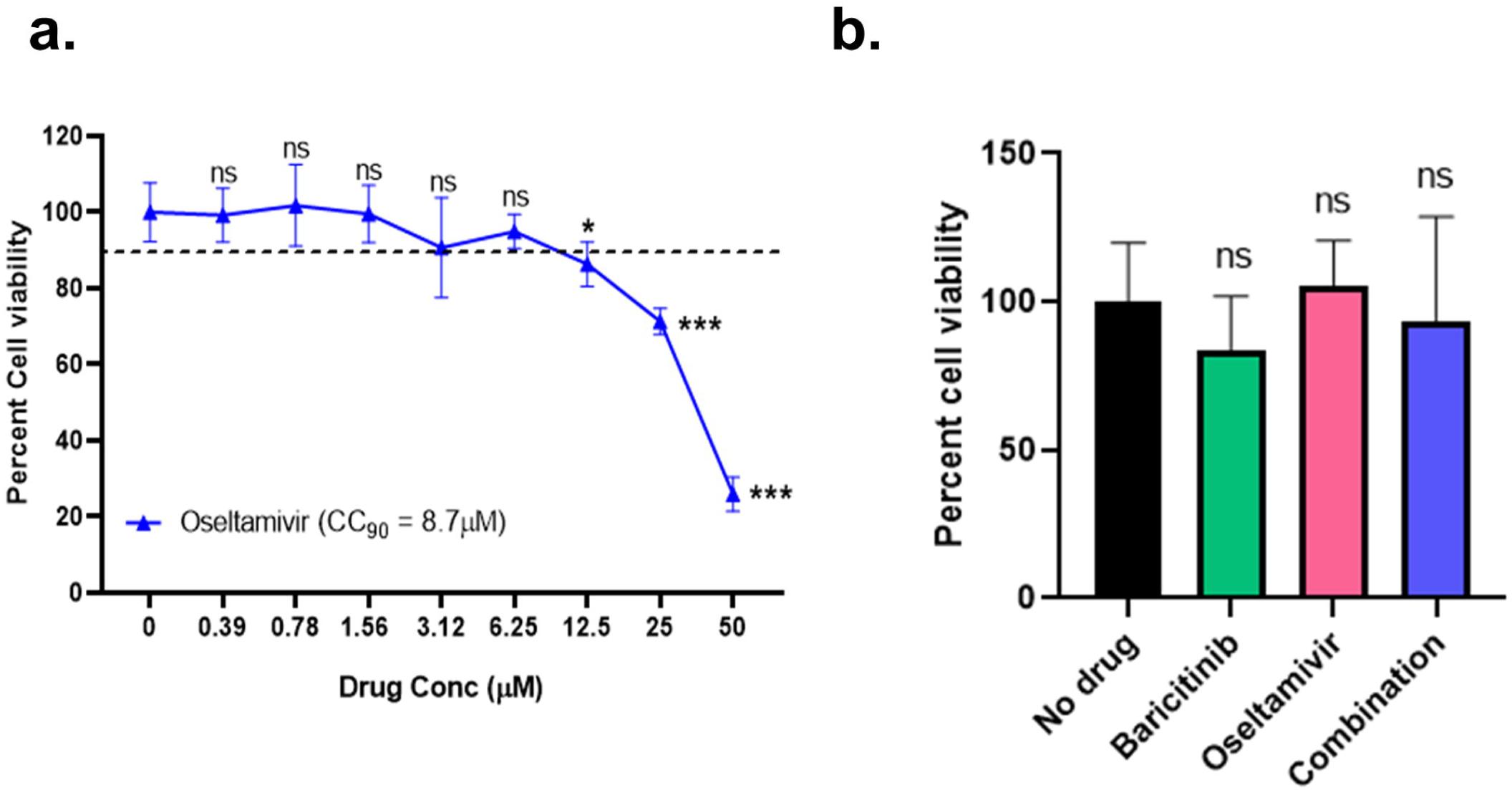

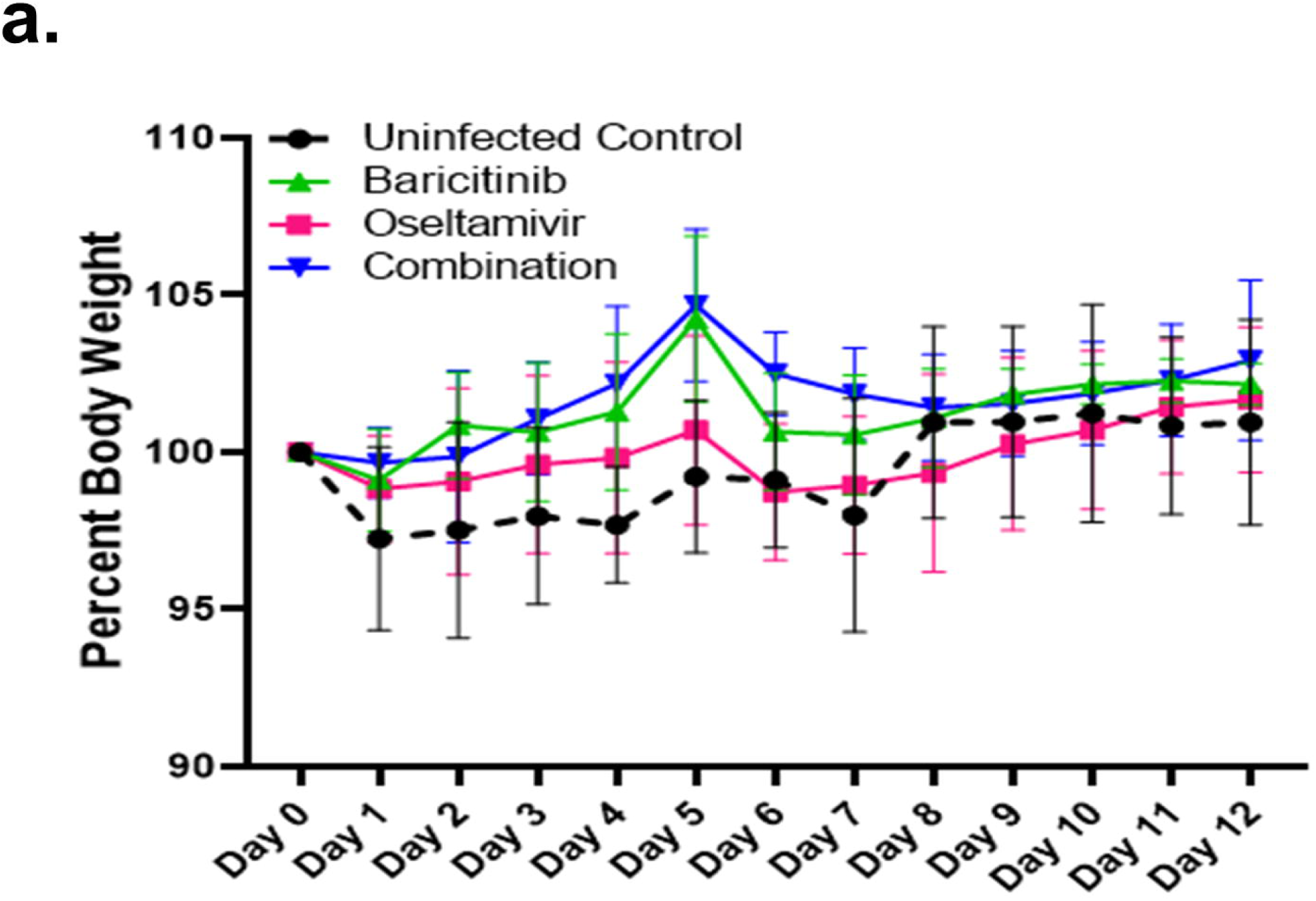

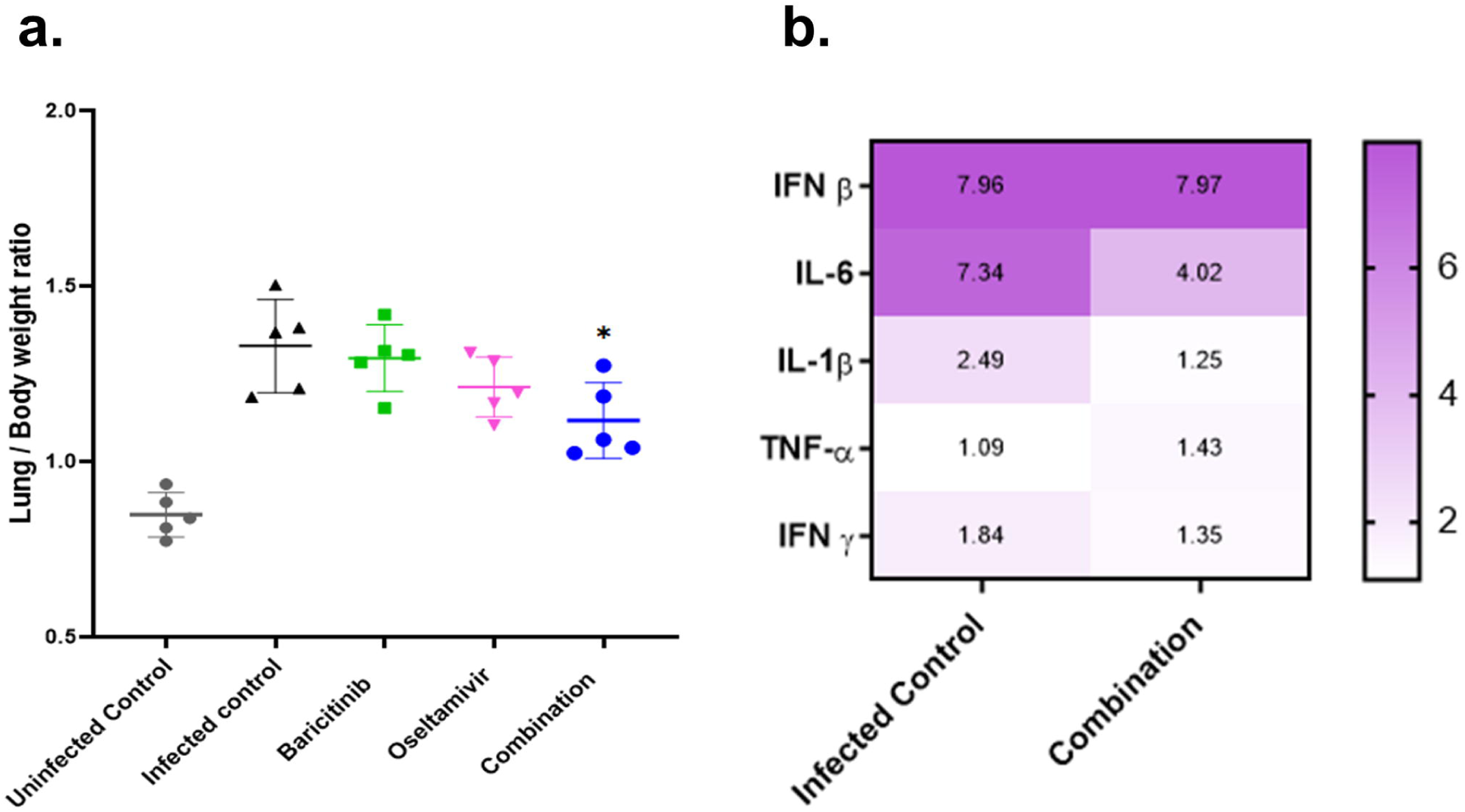

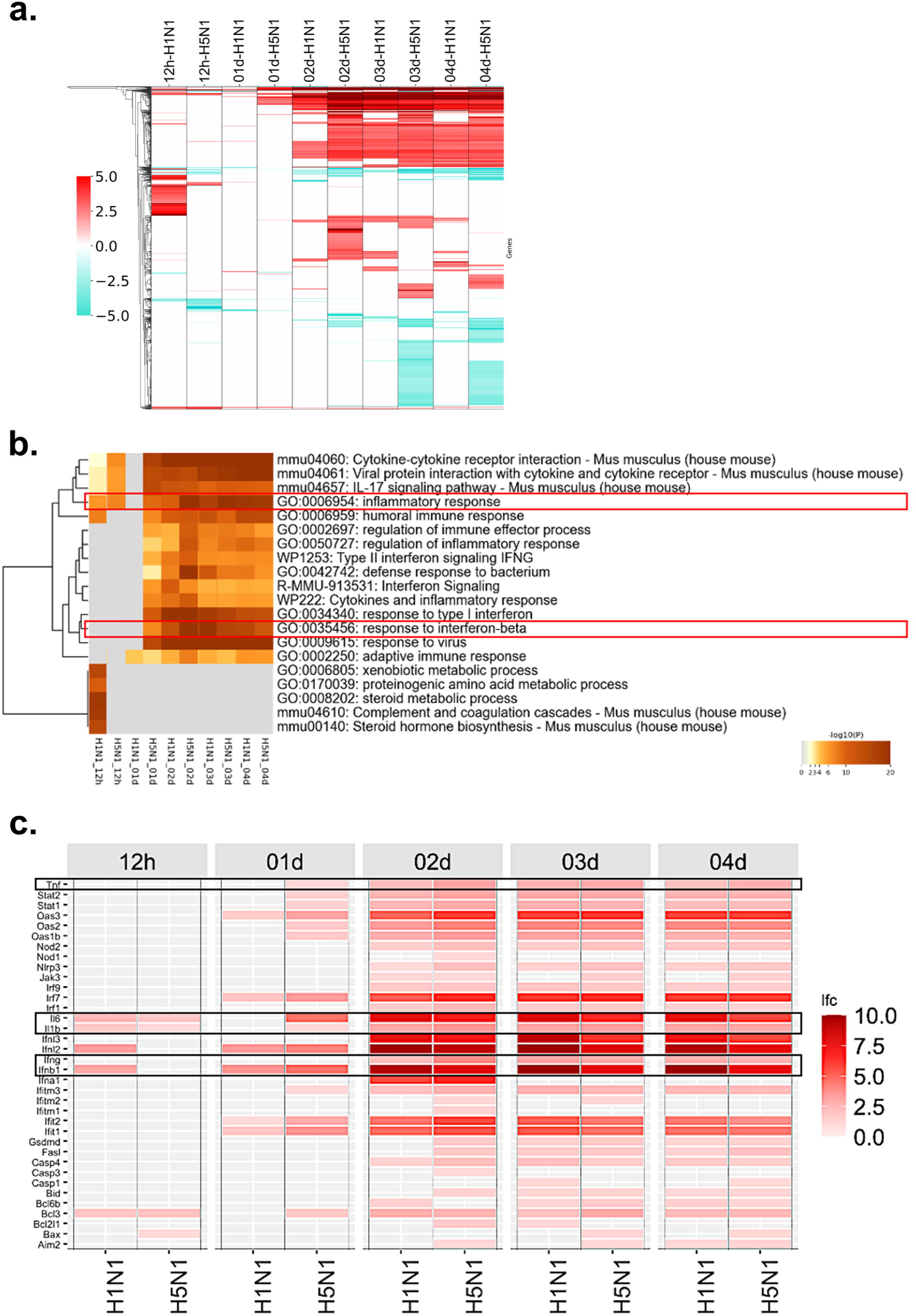

