## Supplementary Data for "Differential modulation of Interferon and Cell Death Responses Define Human vs Avian Influenza A Virus Strain-Specific Virulence and guide Combination Therapy"

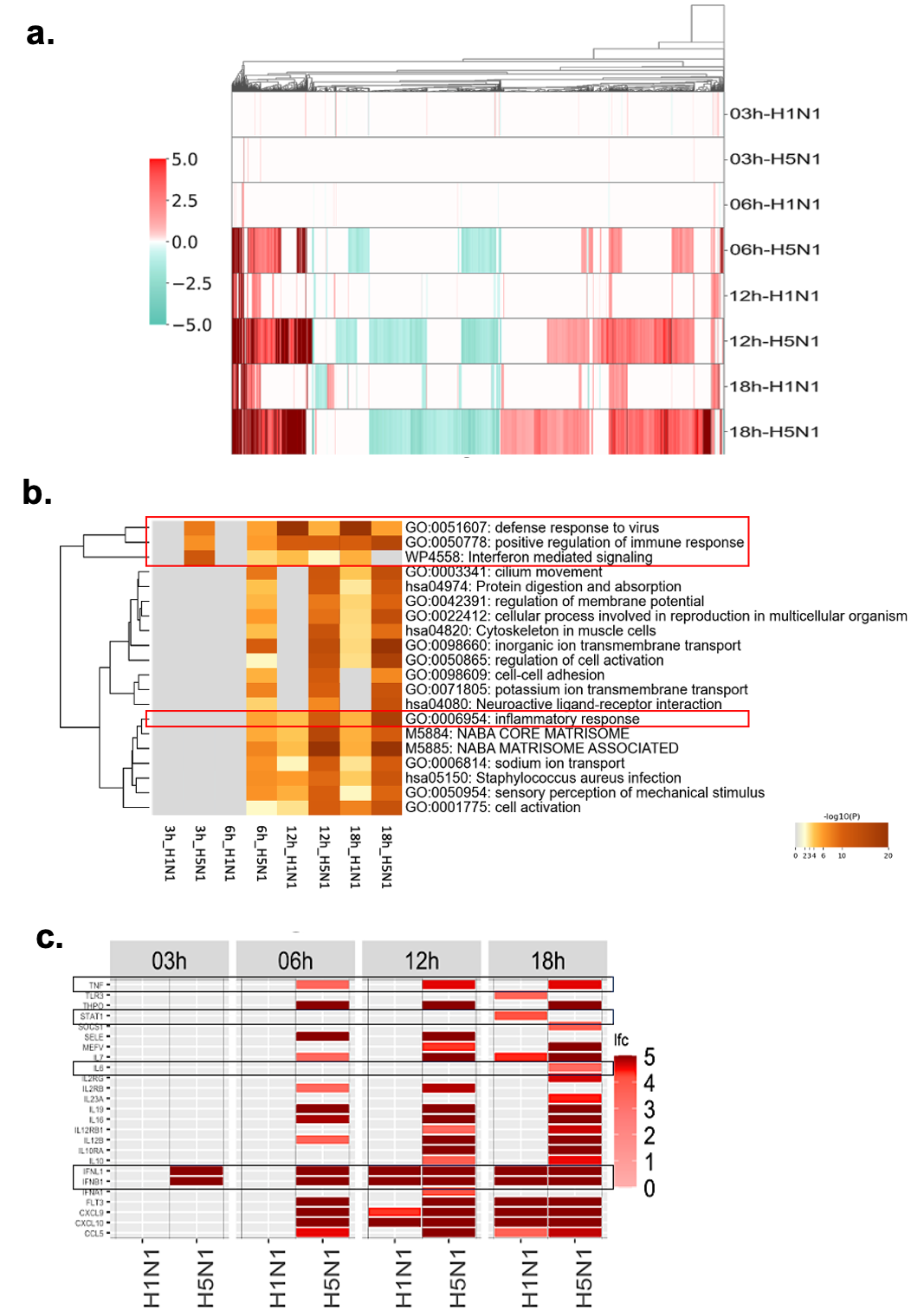

**Supplementary Figure 1. H5N1 and H1N1 IAVs differentially modulate transcription IFN response genes in HTBEs.** Transcriptomics data was obtained from publicly available FluOmics database for differentially expressed genes (DEGs) in HTBE cells upon infection with either H1N1 or H5N1 IAVs 3, 6, 12 and 18 hours post infection **(a)** Cluster map of DEGs was made, where the magnitude of expression is illustrated by the intensity of the colour. **(b)** Heat map of top 20 enriched terms, analyzed by Metascape, coloured by *p* values. (https://metascape.org/gp/index.html#/main/step1). (**c**) The data was further used to determine the temporal expression of differentially regulated genes that code for different cytokines. Cutoff p value ≤ 0.01 and log_2_ fold change (lfc) ≥ 3.32.

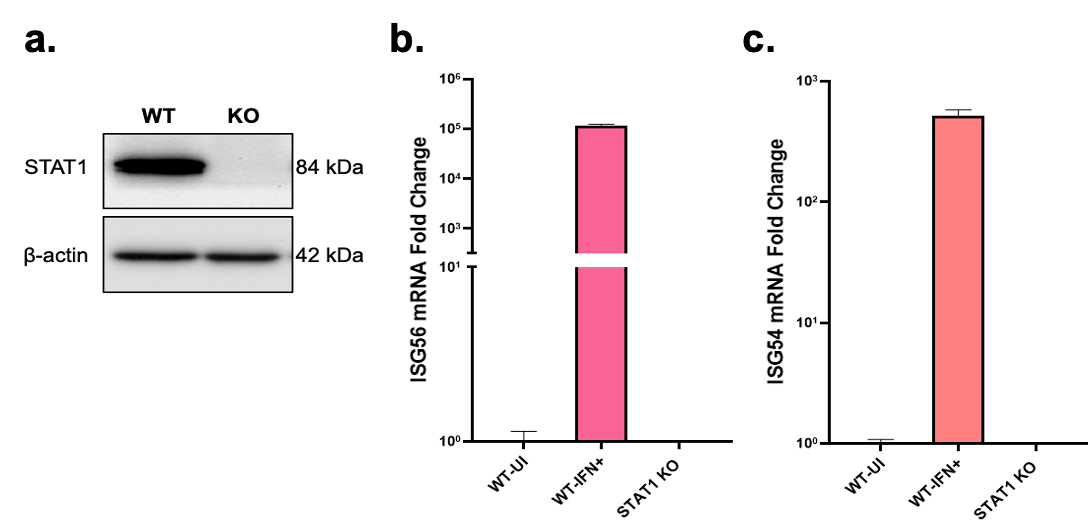

**Supplementary Figure 2. Generation and Validation of A549-STAT1 Knockout Cells. (a)** Cell lysates were analyzed by western blot and probed for STAT1 (84 kDa) and β-actin (42 kDa) **(b)** ISG56/IFIT1 and **(c)** ISG54/IFIT2 mRNA fold change levels were determined by qRT-PCR from total RNA isolated from A549-WT, induced with (1000U/ml) IFN, and A549-STAT1 Knockout cells.

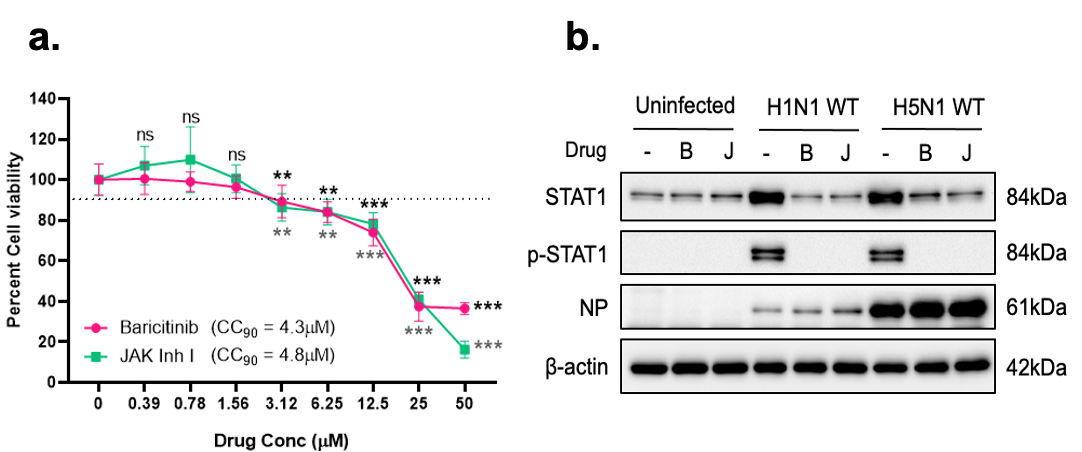

**Supplementary Figure 3**. **Characterization of STAT1 signaling inhibitors (a)** A549 cells were treated with increasing concentration of Baricitinib or JAK Inhibitor I for 48 hours, followed by ATP-dependent luminescence-based cell viability assay to calculate the CC_90_ of the drugs. **(b)** A549 cells were infected with 0.1 MOI of either H1N1 or H5N1 WT IAVs, 6HPI cells were treated with 1 µM of either Baricitinib or JAK Inhibitor I and 48 HPI cells were lysed and inhibition of IFN signaling was analyzed by western blotting for STAT1, pSTAT1, viral NP and β-actin proteins. Data are presented as mean ± SD from triplicate samples of a single experiment and are representative of results from three independent experiments. ∗∗∗, p < 0.001, ∗∗, p < 0.01, and ∗, p < 0.05, by two-way ANOVA mixed effects model with Geisser-Greenhouse correction and Sidak's multiple comparison test (a).

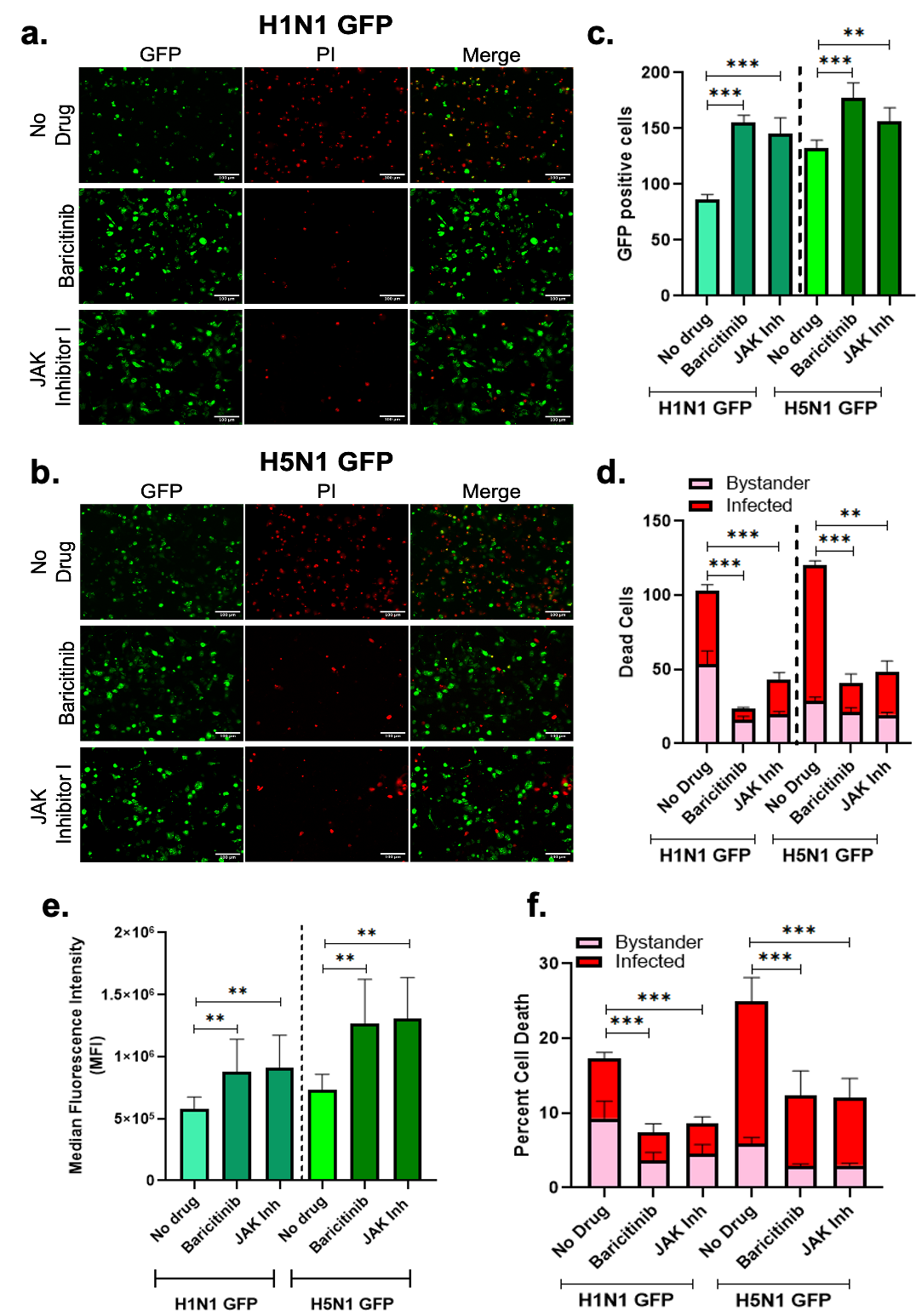

**Supplementary Figure 4. STAT-1 signaling inhibition ameliorates of GFP reporter IAV-induced cell death in human respiratory cells. (a)** A549 cells were infected with either H1N1 or **(b)** H5N1 GFP reporter virus at an MOI of 0.1. 6 HPI cells were treated with 1 µM either Baricitinib or JAK inhibitor I for 48 HPI, the cells were visualized under fluorescent microscope, and the images were quantified for **(c)** GFP-positive (infected) cells; **(d)** cells that were both GFP+PI+ (infected and dead), as well as GFP–PI+ (bystander dead cells), which together represent the total number of dead (PI+) cells. Total GFP+, PI+, and GFP+PI+ cells were counted from six different fields and plotted as cell numbers. **(e)** Similarly, A549 cells were infected with either H1N1 or H5N1 GFP reporter virus at an MOI of 0.1. 6 HPI cells were treated with 1 µM either Baricitinib or JAK inhibitor I for 48 HPI, and the Median Fluorescence Intensity (virual replication) **(f)** cells that were both GFP+PI+ (Q2 infected and dead), as well as GFP–PI+ (Q1 bystander dead cells), which together represent the total number of dead (PI+) cells were quantified using flow cytometry. The data was normalized with respect to the No Drug control. Data are presented as mean ± SD from triplicate samples of a single experiment and are representative of results from three independent experiments. ∗∗∗, p < 0.001, ∗∗, p < 0.01, and ∗, p < 0.05, by Brown-Forsythe and Welch ANOVA testes, Holm-Sidak's multiple comparisions test, with individual variances computed for each comparison. The statistics reflect Total cell death.

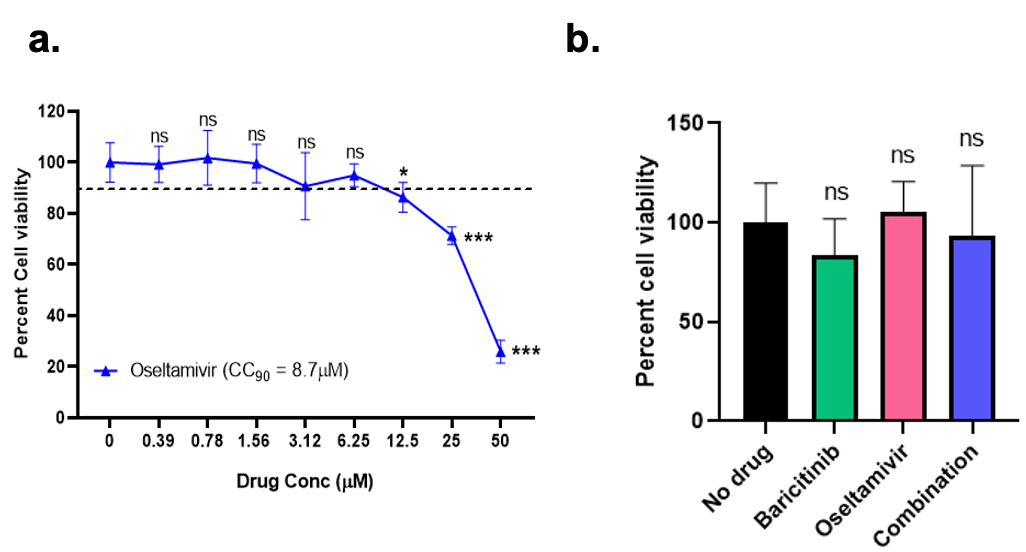

**Supplementary Figure 5: Characterization of Baricitinib and Oseltamivir combination therapy. (a)** A549 cells were treated with increasing concentration of Oseltamivir for 48 hours followed by ATP dependent luminescence-based cell viability assay to calculate CC_90_. **(b)** A549 cells were treated with either Baricitinib, Oseltamivir or their combination till 48 hours to check for cell viability in presence of the combination. Data are presented as mean ± SD from triplicate samples of a single experiment and are representative of results from three independent experiments. ∗∗∗, p < 0.001, ∗∗, p < 0.01, and ∗, p < 0.05, by ordinary one-way ANOVA with Dunnett’s multiple comparisons test.

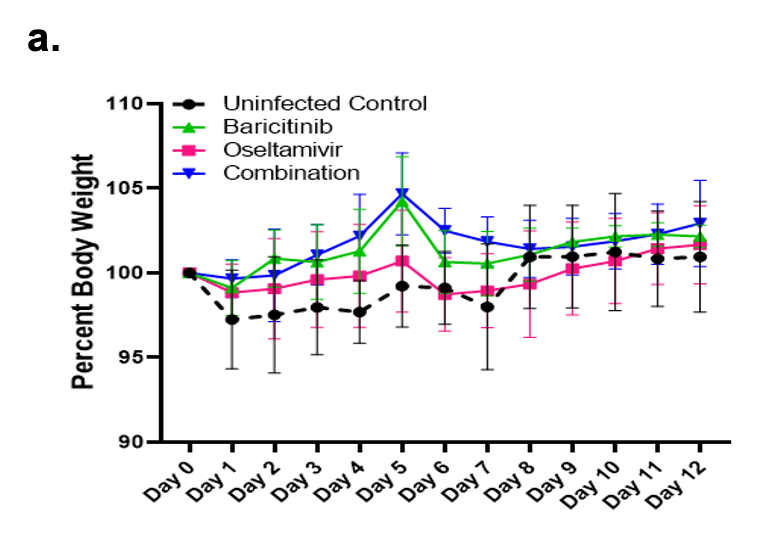

**Supplementary Figure 6**: **Toxicity study of the combination therapy in murine model.** **(a)** Toxicity results as shown by bodyweight changes over 12 days post-treatment with 10 mg/kg of either Baricitinib alone, Oseltamivir alone or their Combination.

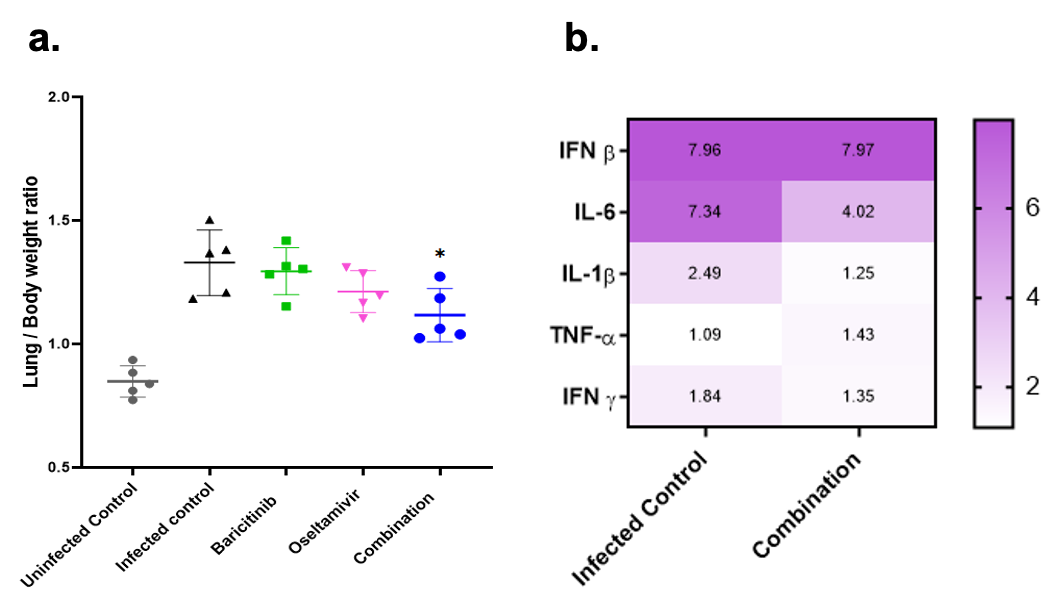

**Supplementary Figure 7: Combination of Baricitinib and Oseltamivir reduces inflammatory gene expression and lung inflammation in H5N1-infected mice. (a**) Lung to body weight ratio at D+4 of infection. **(b)** mRNA fold change (log_2_) levels of several inflammatory cytokines were determined by qRT-PCR from total RNA isolated 4 DPI from lung tissue infected with H1N1 WT IAV and treated with Combination (Baricitinib + Oseltamivir) 2 DPI. Data are presented as mean ± SD from n=5 animals per group ∗∗∗, p < 0.001, ∗∗, p < 0.01, and ∗, p < 0.05, by One-way ANOVA with Dunnett’s multiple comparisons test (a)

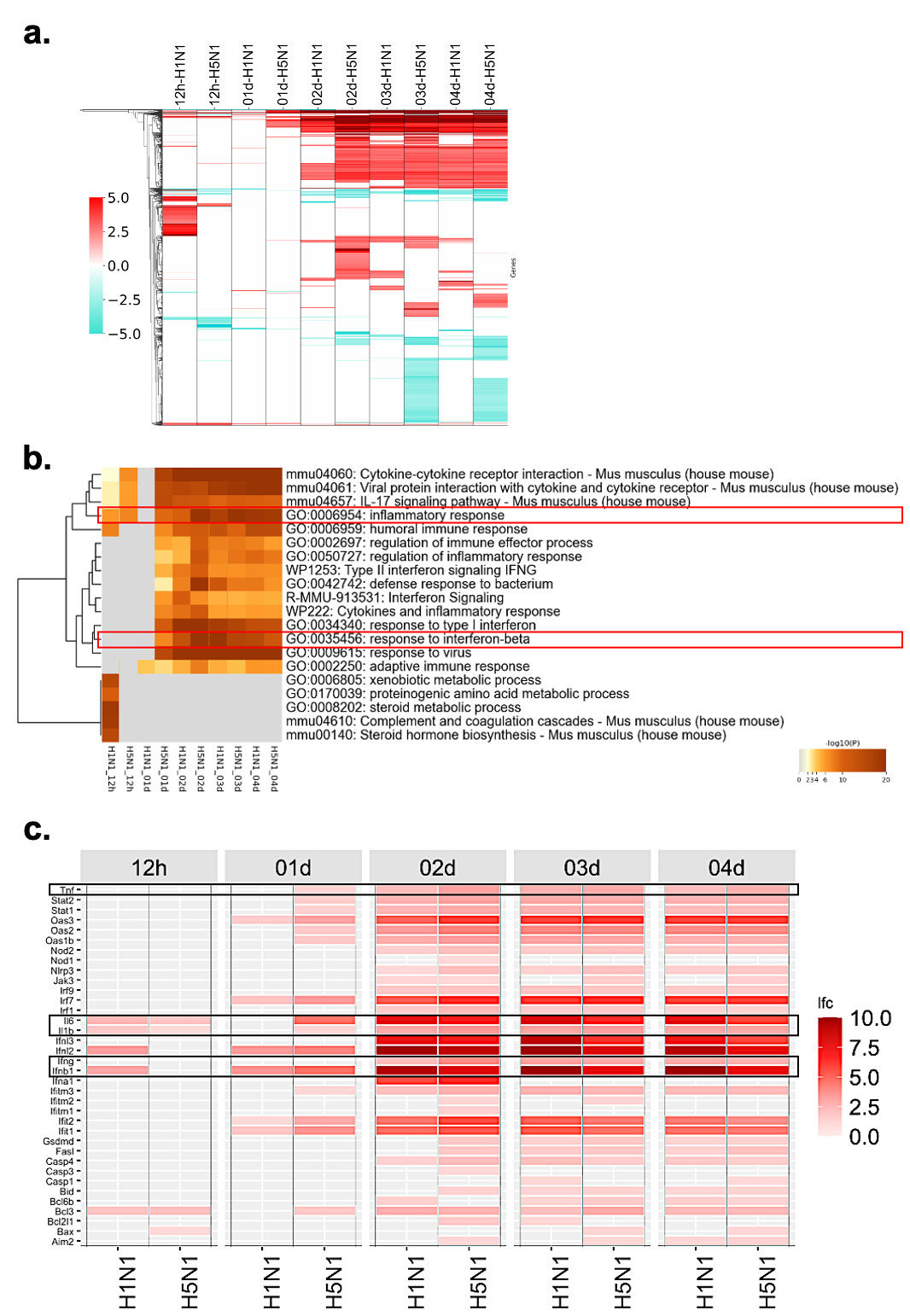

**Supplementary Figure 8: H5N1 and H1N1 IAVs differentially modulate transcription of genes involved in IFN mediated signaling and cell death in murine model.** Transcriptomics data was obtained from publicly available FluOmics database for differentially expressed genes (DEGs) in C57BL/6 mice lung tissue upon infection with either H1N1 or H5N1 IAVs 12 HPI, 1, 2, 3 and 4 DPI **(a)** Cluster map of DEGs was made, where the magnitude of expression is illustrated by the intensity of the colour. **(b)** Heat map of top 20 enriched terms, analyzed by Metascape, coloured by *p* values. (https://metascape.org/gp/index.html#/main/step1). Cutoff p value ≤ 0.01 and log_2_ fold change (lfc) ≥ 3.32. (**c**) The data was further used to determine the temporal expression of differentially regulated genes that code for different cytokines Cutoff p value ≤ 0.05 and log_2_ fold change (lfc) ≥ 1.
